# Innate immunocompetent iNSpheroids: A hiPSC-derived 3D model to study the central nervous system captures an early CNS response to rAAV

**DOI:** 10.1101/2025.07.07.662807

**Authors:** Catarina M Gomes, Gabriela Silva, Mafalda Aleixo, Daniel Simão, Stephan J Holtkamp, Diana D Lobo, Pradeep Harish, Rosalind Jenkins, Lek Dahal, Rui J Nobre, Luís P Almeida, Mark Trautwein, Paula M Alves, Catarina Brito

## Abstract

Gene therapies using adeno-associated viruses (AAVs) for central nervous system (CNS) disorders face challenges due to host immune responses not represented in classical preclinical models. Here, we present a human-induced pluripotent stem cell (hiPSC)-derived innate immunocompetent 3D CNS model that recapitulates neuroinflammatory hallmarks, serving as a platform for preclinical gene therapy development. Utilizing various scales of stirred-tank bioreactor systems, we generated (neurospheroids) iNSpheroids composed of neurons, astrocytes, and oligodendrocytes, alongside microglial cells (iMGL) to mimic the neuro-immune axis. These systems enabled large-scale production of iNSpheroids and subsequent miniaturization for co-culture experiments and screening of inflammatory stimuli, while maintaining a highly controlled environment. The iMGL-iNSpheroids demonstrated active neuron-microglia crosstalk and exhibited distinct inflammatory responses to a series of neuroinflammatory factors. iMGL-iNSpheroids mounted a mild and transient response to rAAV9, mediated by the activation of inflammatory pathways (e.g., TNF-via NF-κB activation) in glial cell populations. This model offers a valuable tool to dissect neuroinflammatory mechanisms, accelerating gene therapy development.

**Teaser:** Immune-competent 3D human CNS model recapitulates glial responses to rAAVs, enabling reliable preclinical gene therapy screening.

## Introduction

Gene therapy using recombinant adeno-associated viral vectors (rAAV) has emerged as a promising strategy for addressing central nervous system (CNS) genetic disorders (reviewed in (*1*)). In rodent and non-human primate (NHP) preclinical models, rAAVs have been shown to sustain gene expression with low immunogenicity risk and genomic integration compared to other viral vectors (*2*). Despite the overall safety and efficacy demonstrated in those CNS-directed rAAV preclinical trials, non-predicted adverse effects have been reported in clinical trials (reviewed in (*3*)). These differences illustrate species-specific variations in the host immune system (*4–7*). Clinical trials for rAAV-mediated gene therapy in the CNS have revealed potential risks associated with the high-dose vector administered (*8*, *9*).High rAAV doses can elicit immune responses and systemic toxicity (*10*). Reports of severe systemic toxicity and fatalities in trials for X-linked myotubular myopathy (MTM) (*11*) and Duchenne muscular dystrophy (DMD) (*12*) underscore the pressing need for a better understanding of host-rAAV interactions, augmenting transduction efficiency, while lowering rAAV dosages. Adverse events are frequently linked to immune responses against the different rAAV components, leading to the activation of cytotoxic T cells and other immune system mediators (*13*). In the CNS, rAAV-induced neuroinflammation, tissue pathology, and behavioural changes have been observed in several NHP models, and recently in clinical studies; astrogliosis and microgliosis have been implicated (reviewed in (*10*)).

Achieving efficient rAAV vector transduction of CNS cells and mitigating immune-mediated adverse effects remain significant challenges. These limitations highlight the need for advanced human in vitro models that can increase our understanding of rAAV biology while improving preclinical accuracy. Using human induced pluripotent stem cell (hiPSC)-based models to elucidate the functional roles of glial cells has the potential to improve advanced therapy development (*14*). Developing immunocompetent human CNS models is essential to better understand neuroimmune interactions and their implications for gene therapy. The emerging three-dimensional (3D) cultured microglia-containing human brain organoids are currently a focus in the CNS modelling field (*15–19*). These in vitro 3D cell cultures recapitulate key features of the human brain, providing a new approach to model brain development and pathology. However, a consensus in the methods used, microglia origin, abundance, and functional state within the organoids is yet to be reached, leading to inconsistent results, particularly in gene and protein expression profiles (*20*).

To address these challenges, in this work, an innate immunocompetent 3D hiPSC-derived CNS model was developed. By integrating major human CNS cell types into a size-controlled, scaffold-free neurospheroid system that enables enhanced reproducibility and immune cell incorporation, our model provides a platform for comprehensively investigating the intricate host-virus relationship, addressing the limitations of traditional organoid-based approaches. We hypothesized that (i) employing neurospheroids differentiated from hiPSC-derived neural progenitor cells (iNSpheroids), which assures consistent and homogenous differentiation into neurons, astrocytes, and oligodendrocytes (*21*), would improve the incorporation of isogenic microglia, the CNS resident immune cells; (ii) and maintaining a controlled microenvironment in computer-controlled stirred tank bioreactors would assure the quiescence of introduced microglia and its functionality. Here, we leverage this model to investigate how microglia shape inflammatory responses and influence rAAV transduction dynamics. Using bulk and single-cell transcriptomics, alongside secretome analysis, we characterized key innate immune pathways activated upon rAAV9 exposure. This model opens new avenues for preclinical research, offering a robust platform to study host-virus interactions and advance the development of safer, more effective rAAV-based gene therapies for CNS disorders.

## Results

### iMGL integrates iNSpheroids retaining identity and acquiring morphoplasticity

Neural culture models incorporating human microglia are essential to advance our understanding of brain cellular interactions and neuroimmune responses under pathological conditions. However, transcriptomic, proteomic, and morphological changes occurring in in vitro cultured microglia present a significant obstacle for human microglia studies (*22*). To develop a physiologically relevant CNS model that more accurately represents neuro-immune interactions, we co-cultured hiPSC-derived microglia (iMGL) with iNSpheroids - composed of neuronal and macroglial cells (*23*, *24*). We hypothesized that employing a size-controlled neurospheroid system in a physiological environment would ensure direct microglia incorporation. This setup would promote physiologically relevant cell-cell interactions and surpass the limitations of conventional organoid systems, which often lack the integration and maturation of microglia. iMGL were differentiated from two hiPSC lines using a two-stage protocol (*17*): (i) erythro-myeloid progenitor (iEMP) differentiation from hiPSCs (12 days) and (ii) iEMP differentiation into iMGL (28 days) (Fig. S1 A). Over 80% of iEMPs (D12) expressed the hematopoietic progenitor markers CD43 and CD45 while exhibiting low expression of pluripotent (TRA-1-60, TRA-1-81, and SSEA4) and microglial (CX3CR1 and TREM2) lineage markers (Fig. S1 B, C). By the end of the 40-day protocol, iMGL expressed the lineage markers, IBA-1 and TREM-2 (Fig. S1 B, C). These cells differentially responded to prototypical inflammatory stimuli, acquiring distinct morphological profiles (Fig. S1 D). This was particularly evident for the LPS + IFN-γ treatment, where most cells acquired a semi-adherent and round morphology (Fig. S1 D). This differential response was also evident at the gene expression level, with the significant upregulation of immunomodulators, namely CXCL8, 24h hours upon stimuli such as TNF-α, IL-1α and C1q (collectively known as TIC) or LPS plus IFN-γ (Fig. S1 E). The anti-inflammatory stimulus of interleukin(IL)-4 plus IL-13, had a low effect on the target genes evaluated, with some apparent downregulation at 6 hours post-stimulus.

iNSpheroids were generated by differentiating hiPSC-NPC spheroids (neurospheres) into iNSpheroids in perfusion stirred-tank bioreactors (STBs) (*23–26*). iNSpheroids recapitulate features of the brain microenvironment, such as deposition of a human brain-like extracellular matrix, and containing more mature neurons and astrocytes in comparison to other 3D models, establishing neuron-glia interactions such as functional glutamine-glutamate-GABA shuttle (*21*, *24*). Two commercially available hiPSC lines were employed for iNSpheroid differentiation: (i) iPSC(IMR90)-clone 4, as reported previously (*26*), and iPSC(DF6-9-9T.B), (Fig. S2). The two cell lines presented similar neurosphere formation kinetics, spheroid morphology, and diameter throughout time (Fig. S2 A, B). Similarly to what was previously reported for iNSpheroids derived from iPSC(IMR90)-clone 4 (*26*), the neural stem cell phenotype was maintained during the first 7 days in culture for the iPSC(DF6-9-9T.B) line (Fig. S2 C). Likewise, proliferation arrest (shown by PCNA downregulation, Fig. S2 D) and the expression of gene and protein markers for neurons (Fig. S2 E, F), astrocytes (Fig. S2 G), and oligodendrocytes (Fig. S2 H) further supported the lineage commitment towards neural differentiation.

Differentiated iMGL and iNSpheroids were co-cultured in the Ambr® 15 Cell Culture STB (Figure 1 A), to attain a controlled environment. To characterize the iMGL-iNSpheroid phenotype in detail, sn-RNAseq analysis of 3-day co-cultures was performed. Assessment of cell type composition (clustering and annotation, Fig. 1 B and Fig. S3) revealed the presence of neuronal, astrocytic, oligodendrocyte, and microglial populations (Fig. 1 B and Fig. S3 A, B). Cells in cluster 5, annotated as microglia, expressed microglia typical genes, such as AIF1, SPP1, CSF1, P2RY12 and −13, CX3CR1, HLA-DRA and GPR34 (Fig. 1 C and Fig. S3 B). The expression of such genes was further evaluated by bulk RNAseq, emphasizing the expression of numerous human microglia homeostatic (e.g., TMEM119, P2RY12) and function-related genes, namely inflammatory machinery (e.g., interferon and MHC genes) (Fig. 1 D). Immunolabelling of microglial proteins corroborated the successful incorporation and maintenance of iMGL within the 3D structure of the iNSpheroids (Fig. 2 A). Detection of IBA-1, TMEM119, and P2RY12 confirmed microglia identity (Fig. 2 A-C). Notably, viable iMGL were also observed in the supernatant throughout the ten days of co-culture (Fig. S4 A, white-outlined square). Within the iNSpheroid 3D structure, iMGL with distinct morphological traits were observed (Fig. 2 D-I). Quantitative analysis performed in iMGL across different iNSpheroids revealed a negative correlation between cell circularity and perimeter (Fig. 2 D, r2 = −0.90), and a positive correlation between cell length and perimeter (Fig. 2 E, r2 = 0.82). The average iMGL, area was 150-200 μm2 (Fig. 2 F). This iMGL, area was close to the lower limit of the physiological range of 200–8000 μm2 in the human brain (*27*). iMGL, cell bodies were found in different Z-planes (depth >35 μm, Fig. 2 G) and branches extending at >20 μm in optical depth (Fig. 2 H and I), within iNSpheroids, of approximately 100-200 μm, of diameter (Fig. S2 B). Quantifying ramification length in the iMGL-iNSpheroid structure is challenging given their organization in different planes and depths of the iNSpheroid (Fig. 2 H and I). Still, by observing iMGL, branching, we could identify ramifications extending up to 30 μm (Fig. 2 G-I). Altogether, the data suggests that iMGL, extended processes to sense the iNSpheroid microenvironment in distinct Z-planes, suggesting a scavenging function as reported for microglia in the healthy brain parenchyma (*27*, *28*).

**Fig. 1.**
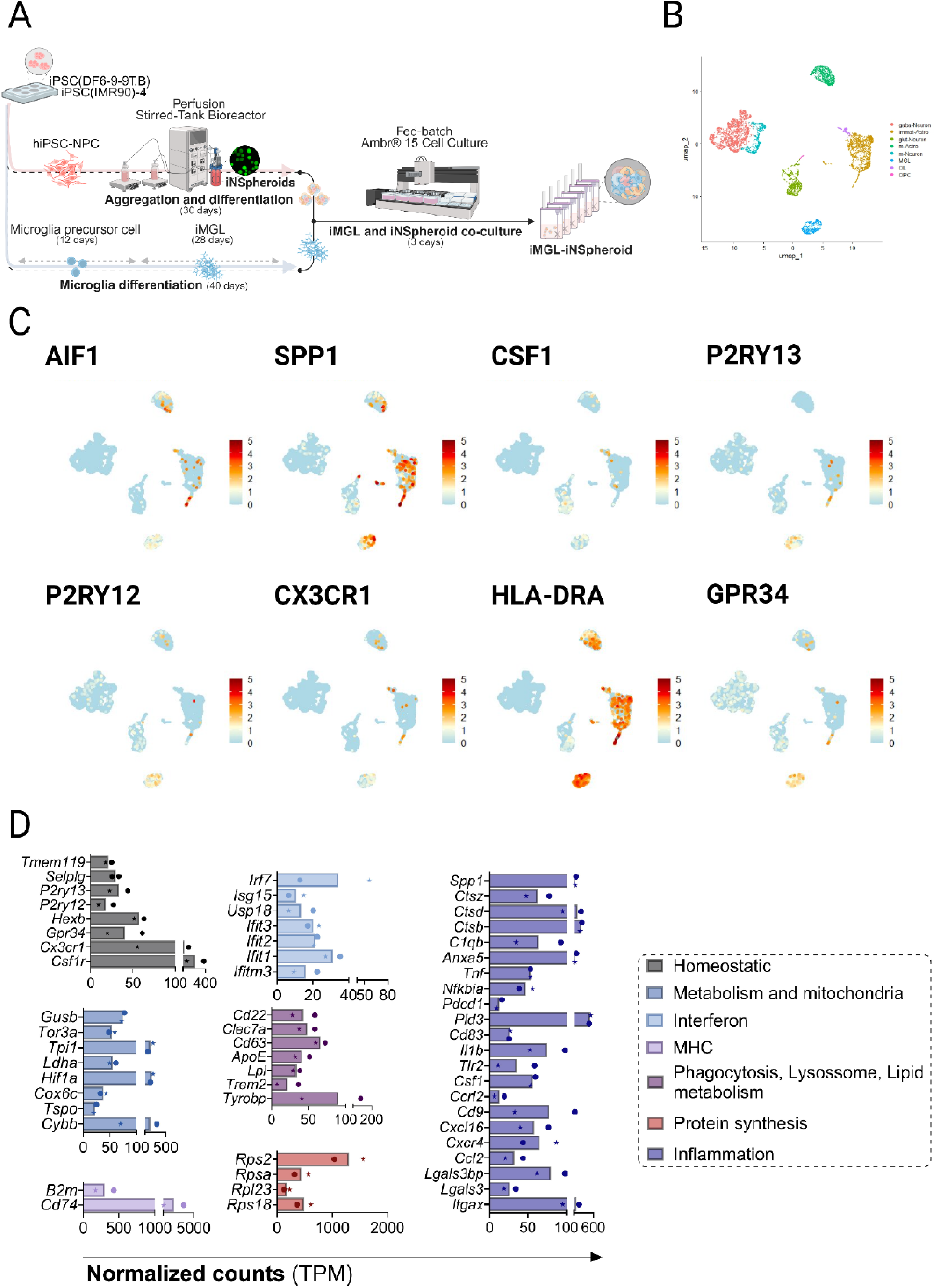
**iMGL within iMGL-iNSpheroids retains cell identity, after three days of co-culture.** (a) Schematic representation of the parallel differentiation of human induced pluripotent stem cell (hiPSC)-derived microglia (iMGL) and hiPSC-derived neurospheroids (iNSpheroids) and the strategy to attain their co-culture (iMGL-iNSpheroids). The experimental design involves the differentiation of iNSpheroids concomitantly with iMGL from the same hiPSC line to obtain isogenic co-cultures. The iNSpheroids were obtained in 250 mL bioreactors and transferred to a microbioreactor system (15 mL) for co-culture. (b) UMAP plots of single nuclei RNA-sequencing (sn-RNAseq) generated with Seurat. Clustering the integrated datasets at resolution 0.1 allowed identifying seven major clusters, corresponding to the major neural cell types (neurons, astrocytes, and oligodendrocytes) and microglia. UMAP projection calculated with the first 30 PCs. (c) UMAP plots of sn-RNAseq generated with Seurat. Cells are clustered in two dimensions using the UMAP dimensionality reduction methodology and annotated by gene expression. Each coloured dot represents a cell expressing the represented gene, while the colour code represents the gene expression level. The data corresponds to two integrated datasets, from two hiPSC lines (iPSC(IMR90)-clone 4 – circle – and iPSC(DF6-9-9T.B) – star symbol). (d) RNA-Seq expression levels in transcripts per million (TPM) of microglial genes related to homeostasis, metabolism, protein synthesis, interferon- and MHC-pathways, phagocytosis, and inflammation. Dots correspond to the two hiPSC lines used in this study. Data represented as mean ± standard deviation (S.D.) (N = 2).

**Fig. 2.**
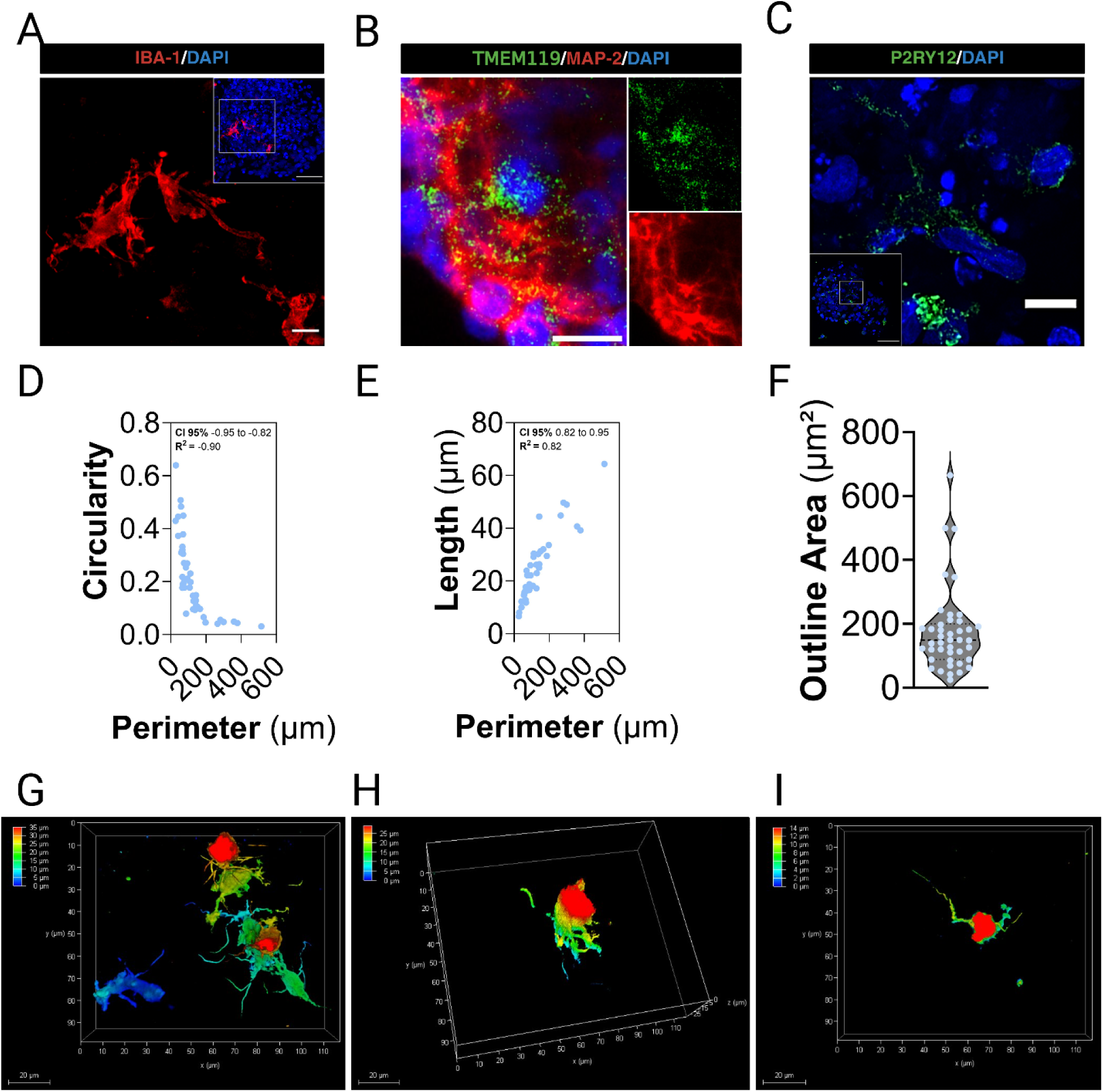
**iMGL within iMGL-iNSpheroids expresses canonical markers and exhibit morphological plasticity.** (a) Confocal images showing IBA-1 (red), (b) TMEM119 (green) and MAP-2 (red), and (c) P2RY12 (green) immunolabelling, all counterstained with DAPI, indicating the presence of microglia within the iNSpheroid 3D structure. Scale bars: 50 and 10 μm. Correlation analysis between: (d) cell perimeter and circularity; (e) cell perimeter and branch length. (f) Quantitative analysis of cell outline area. Each dot corresponds to one IBA-1-positive cell, data from 5 different iNSpheroids. (g) 3D reconstruction of confocal images of IBA-1 labelling, illustrating microglial morphological complexity, (h) branching in different planes, and (i) branch length. Scale bars: 20 μm.

### iMGL presence improves neuronal synaptic activity in iNSpheroids

The complex range of microglia morphologies and functions in the healthy brain is influenced by the structure of the neuronal networks and activity (*29*). Functional integration of iMGL and its impact on neuron and synapse maturation were studied employing super-resolution acquisition mode in confocal microscopy. Ameboid-like iMGL cells were detected within the iNSpheroids in a homeostatic-like environment (Fig. 3 A), expressing CD68, a myeloid cell lineage glycoprotein associated with phagolysosomes. Fragments of nuclei (DAPI-positive) and dendrites (MAP2-positive) were found within the late endosomes/ phagolysosome-like structures positive for CD68 (Fig. 3 A).

**Fig. 3.**
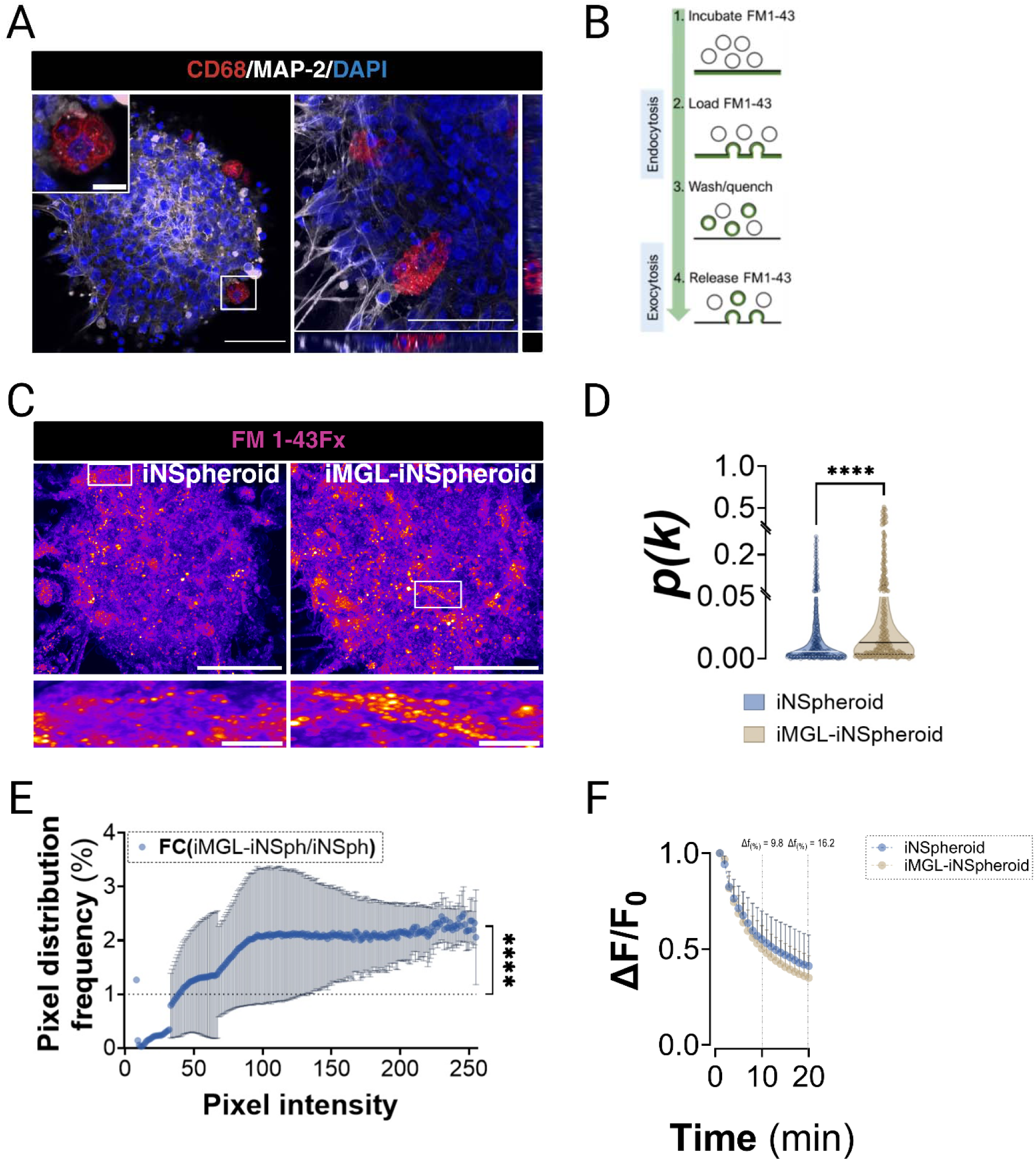
**Microglia interact with neuronal populations within iNSpheroids.** Immunostaining of (a) CD68(red)-positive amoeboid microglia within the 3D MAP2(grey)-positive neuronal network. Nuclei are counterstained with DAPI (blue). The inset highlights microglial and neuronal interactions through phagocytic-like processes. Scale bars: 50 and 10 μm. (b) Schematic representation of the endocytosis and exocytosis assay used to analyse neuronal functionality in iNSpheroids, with and without iMGL. (c) Confocal microscopy of FM 1-43Fx-loaded synaptic vesicles in the iNSpheroids with and without iMGL, demonstrating endocytosis. FM 1-43Fx staining represented using the FIRE function in ImageJ (FIJI) v1.54j. Scale bars: 50 and 10 μm. Quantitative analysis of (d) probability of occurrence of the different intensity values in each pixel and (e) pixel distribution versus intensity as a fold change of the iMGL-iNSpheroid/ iNSpheroid. (f) Fluorescence decay over 20 minutes of the FM 1-43-labelled synaptic vesicles, upon stimulation with a high potassium solution, to elicit synaptic vesicle release. Statistical analysis: One-way ANOVA, ****p< 0.0001. Data are presented as mean ± S.D of N = 4 from two hiPSC lines (iPSC(IMR90)-clone 4 and iPSC(DF6-9-9T.B)).

In previous reports, we have shown that differentiated iNSpheroids can contain negligible cell death(*23*). The presence of these cells may be sufficient to trigger cell debris removal mechanisms (phagocytosis) and synaptic pruning in microglia (reviewed in (*30*, *31*)). To assess the impact of microglia activity on neuronal functionality within the iNSpheroids, we evaluated synaptic vesicle content and dynamics (Fig. 3 B). Synaptic vesicle trafficking (endocytosis and exocytosis) was assessed with the fluorescent probe FM™ 1-43 (*32*). iNSpheroids with or without iMGL were loaded with an FM™ 1-43 analogue (FM™ 1-43FX) to identify synaptic vesicle formation upon extracellular quenching.

Synaptic vesicle-like particles were observed in iNSpheroids with or without iMGL (Fig. 3 C). However, iMGL-iNSpheroids showed a higher distribution of pixels with high fluorescence intensity (Fig. 3 C-E). Applying a neuron-specific depolarizing stimulus evoking vesicle release, iMGL-iNSpheroids showed a tendency for quicker loss of fluorescence compared to iNSpheroids alone (Fig. 3 F), with a decay of 9.8 and 16% higher than the iNSpheroids alone, after 10 and 20 min, respectively. The improvement in both synaptic vesicle formation and release suggests a contribution of iMGL to attain higher neuron maturation, with functional synaptic terminals able to fuse the synaptic vesicles with the plasma membrane and perform endocytosis, exocytosis, and neurotransmitter release.

Given the astrocytic and microglial ability to perform phagocytosis of synapses, apoptotic neurons, and degenerating axons, we next sought to investigate the expression distribution of phagocytosis-related genes in the different cell clusters. Transcriptome data analysis revealed the expression of phagocytosis/ late endosome/ lysosome gene markers, namely CD68, TREM2, CLEC7A, and TYROBP (Fig. 4 A), which in general were identified in the microglia annotated cluster. Complement components C1q and C3, produced by microglia and astrocytes, have been shown to bind to altered neuronal surfaces, signaling these damaged areas for phagocytosis by microglia (*33*). In iMGL-iNSpheroids, we identified puncta positive for both C3 and C1Q in MAP2-positive neurons (Fig. 4 B-D).

**Fig. 4.**
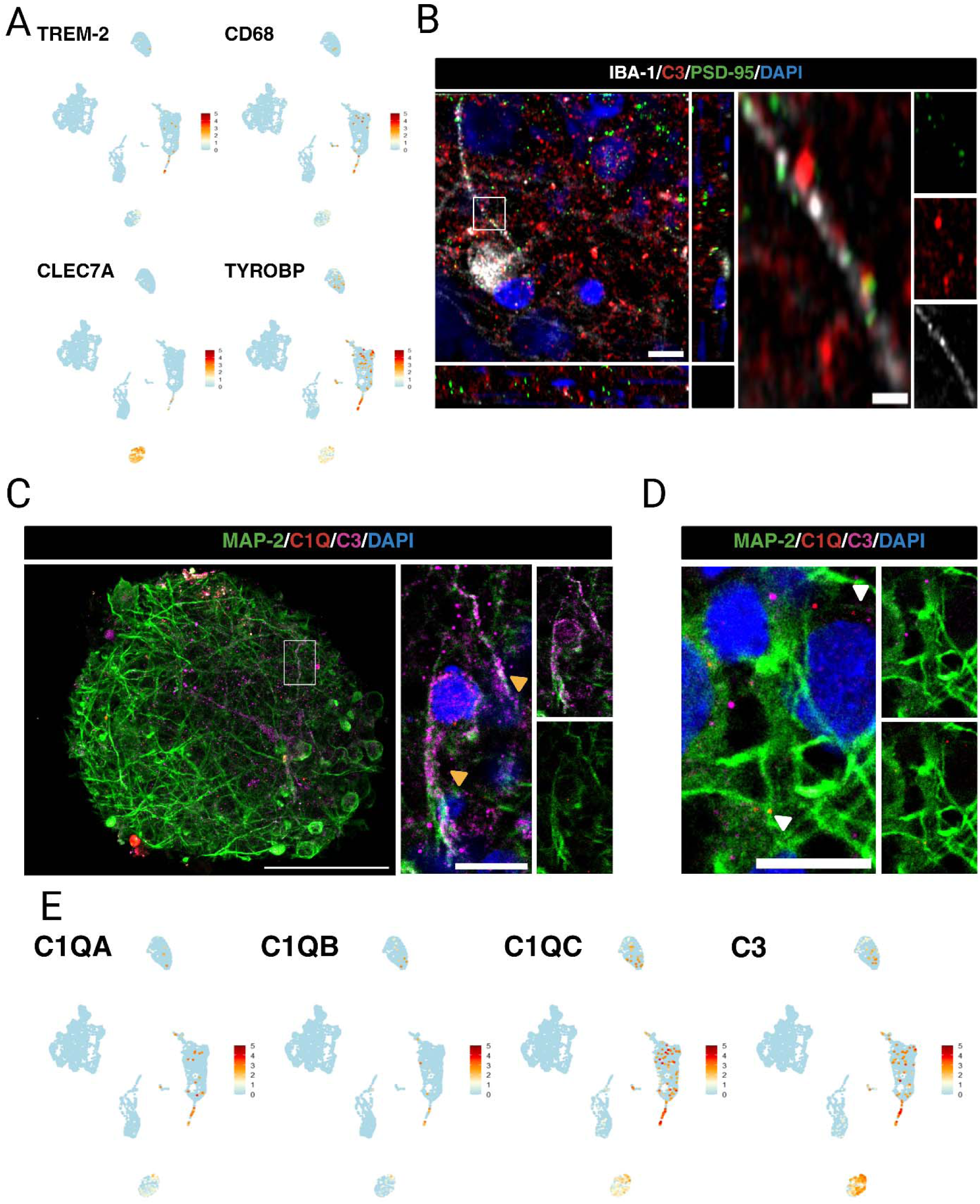
**iMGL within the iNSpheroids presents phagocytic machinery.** (a) UMAP plots of single nuclei RNA-sequencing (sn-RNAseq) generated with Seurat. Cells are clustered in two dimensions using the UMAP dimensionality reduction methodology and annotated by gene expression. Each coloured dot represents a cell, and the colour (ranging from light blue to red) represents the expression level for each gene represented. The data corresponds to two integrated datasets, from two hiPSC lines (iPSC(IMR90)-clone 4 and iPSC(DF6-9-9T.B)). (b) Immunostaining of IBA-1 (grey), PSD-95 (green), and C3 (red), and counterstaining of nuclei with DAPI (blue), showing microglia in close proximity to synaptic structures within iMGL-iNSpheroids. Insets provide higher magnification of the microglial-synaptic interaction. Scale bars: 5 and 2 μm. (c, d) Immunostaining of MAP-2 (green), C1Q (red), and C3 (magenta), and nuclei counterstained with DAPI (blue), showing neurons expressing complement component proteins. Scale bars: 50 and 10 μm. (e) UMAP plots of sn-RNAseq generated with Seurat. Cells are clustered in two dimensions using the UMAP dimensionality reduction methodology and annotated by gene expression. Genes represented: C1QA, C1QB, C1QB and C3. The data corresponds to two integrated datasets, from two hiPSC lines (iPSC(IMR90)-clone 4 and iPSC(DF6-9-9T.B)).

IBA-1-positive microglia was found in close proximity to PSD-95- and C3-double positive puncta, which suggests that microglia-neuron crosstalk can occur through this mechanism. Concomitantly, C3, C1QA, C1QB and C1QC were expressed mainly in the microglia and astrocytic populations (Fig. 4 E).

### Prototypical inflammatory stimuli induce distinct inflammatory signatures in iNSpheroids with or without iMGL

To assess the iMGL-iNSpheroid capacity to mount an immune response, two different proinflammatory stimuli were used (Fig. 5 A): (i) TNF-α, IL-1α, and C1Q (TIC), and (ii) lipopolysaccharide and interferon-gamma (LPS plus IFN-γ). These cocktails are known to be produced during pathogen infection (*34*) and/or neurological disorders, such as Alzheimer’s disease (AD), in patients and mouse models (*35*, *36*). To mimic a neuroprotective environment, a combination of IL-4 and IL-13 was also used. These immunomodulators have been linked to an anti-inflammatory effect in various CNS pathologies, such as multiple sclerosis, AD, and autoimmune encephalomyelitis (*37*). Cell viability was maintained in non-treated and treated conditions, in both iMGL integrated into iNSpheroids and in suspension (Fig. S4 A, B).

**Fig. 5.**
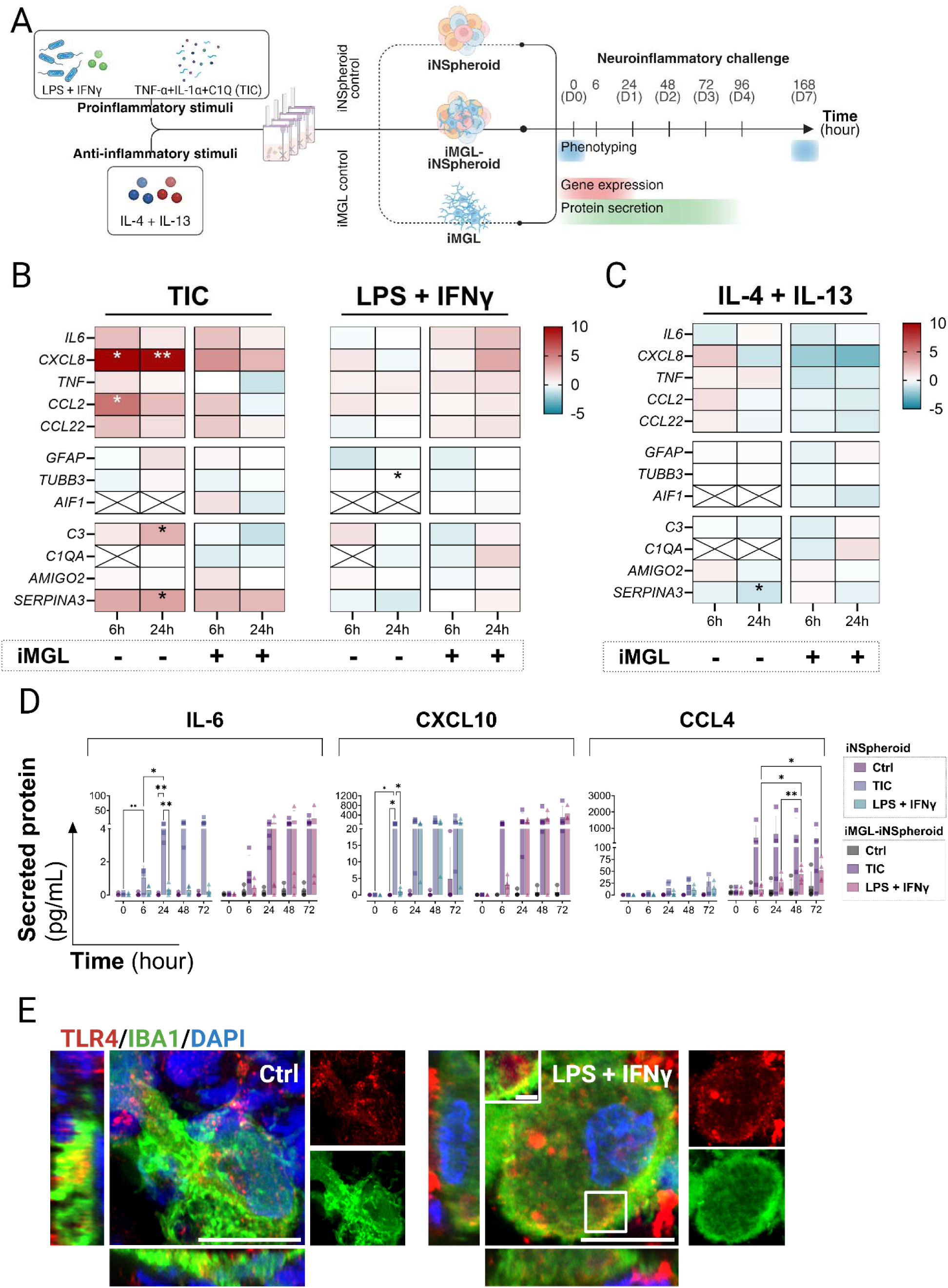
**Prototypical inflammatory stimuli elicit distinct neuroinflammatory responses in iNSpheroids with or without iMGL.** (a) Schematic overview of experimental setup. iNSpheroids were cultured alone or co-cultured with iMGL (iMGL-iNSpheroid) for three days in a microbioreactor system. Cultures were subjected to neuroinflammatory challenges using prototypical inflammatory stimuli: proinflammatory: LPS + IFN-y, TNF-a + IL-1a + C1Q (TIC) or anti-inflammatory: IL-4 + IL-13. Samples were collected at different time points for phenotyping, gene expression (0-, 6-, and 24-hours post-stimuli), and protein secretion (0, 24-, 48-, 72- and 96-hours post-stimuli) analyses. (b, c) Gene modulation quantified for (b) proinflammatory (TIC and LPS + IFN-y) and (c) anti-inflammatory (IL-4 + IL-13) conditions and represented as heatmaps showing Log2 fold changes in gene expression across different conditions and timepoints over non-stimulated controls. Expression was measured for neuroinflammatory (IL-6, CXCL8, TNF-a, CCL2, CCL22), astrocytic (GFAP), reactive astrocytes (C3, C1QA, AMIGO2 and SERPINA3) neuronal (TUBB3), and microglial (AIF-1) genes, in iNSpheroid alone (-), and iMGL-iNSpheroids (+) conditions. Data were generated using RT-qPCR and the comparative cycle threshold value method (2 −ΔΔCt) and are represented as the mean of five independent experiments. Statistical analysis was performed by applying the one-way ANOVA test, *p < 0.05, **p < 0.01. (d) Quantification of secreted IL-6, CXCL10, and CCL4 proteins, identified by secretome analysis at 0, 6, 24, 48, and 72 hours, in iNSpheroids and iMGL-iNSpheroids, stimulated with prototypical stimuli. Immunostaining of toll-like receptor (TLR)4 (red) and IBA-1-positive microglia (green) in (e) iMGL-iNSpheroids.

iNSpheroids with and without iMGL presented different inflammatory profiles in response to the stimuli (Fig. 5 B-D). Several inflammatory modulators, such as IL-6 and CXCL8, were upregulated at 6 and 24 hours upon pro-inflammatory stimulation (Fig. 5 B), in comparison to the unstimulated control. Concomitantly, a tendency for higher secretion of cytokines and chemokines, namely IL-6, CCL4 and CXCL10, was detected (Fig. 5 D, Fig. S5 A, B). iNSpheroids alone responded mostly to the TIC stimulus, by upregulating several of the inflammatory genes analysed, namely CXCL8 and C3. Notably, C3 gene expression increased from 6 to 24 hours post-TIC stimulus (Fig. 5 B). However, the presence of iMGL in iNSpheroids abolished these differences. C3 has been considered a hallmark of astrocyte-microglia crosstalk, being secreted and uptaken by these cells governing their inflammatory loop (*35*). The low expression of toll-like receptor (TLR)4 in human astrocytes, which recognizes LPS at the cell membrane (*38*, *39*), supports the lack of inflammatory response in the iNSpheroids to the LPS plus IFN-γ treatment. Indeed, TLR4 was detected in juxtaposition with IBA1-positive iMGL (Fig. 5 E) and within vesicular-like structures in iMGL cytoplasm (Fig. 5 E, white-outlined square).

Under the anti-inflammatory stimulus, iNSpheroids presented a tendency for upregulation of genes encoding chemokines such as CXCL8, CCL22, CCL2, and TNF, after 6 hours of stimulation (Fig. 5 C). Residual concentration of CXCL8 has been shown to promote cytoprotective effects (*40*, *41*), lead to the production of growth factors (*42*), and modulating synaptic transmission in brain cells (*43*). Still, minimal to no differences were found at the secreted protein level (Fig. S5 C). On the other hand, iMGL-iNSpheroids presented a general downregulation of all inflammation-related genes analysed (Fig. 5 C), suggesting a transcriptional remodelling towards a neuroprotective state, as reported previously (*37*, *44*). Likewise, the secretion of cytokines and chemokines did not increase in comparison to the unstimulated control (Fig. S5 D). Overall, the presence of iMGL impacted the gene expression of the inflammatory modulators analysed, demonstrating the model’s innate immunocompetence.

### rAAV9 transduces iNSpheroids with and without iMGL, with tropism towards neurons and astrocytes

Given the limited literature on the transduction of rAAV9 in human CNS models containing microglia, we first sought to characterize the ability of this viral vector to transduce iNSpheroids. iNSpheroids with and without iMGL were transduced at 5 x 105 VG/cell with three different rAAV9 constructs (Table S3). The multiplicity of infection (MOI (VG/cell)) was selected based on previous reports (*26*), to attain significant transduction efficiencies while avoiding vector-induced toxicity (*45*, *46*). iNSpheroids with and without iMGL were permissive to rAAV9 transduction with similar transduction efficiency and negligible cell death for all the constructs tested (Fig. 6 A-C and Fig. S6 A, B).

**Fig. 6.**
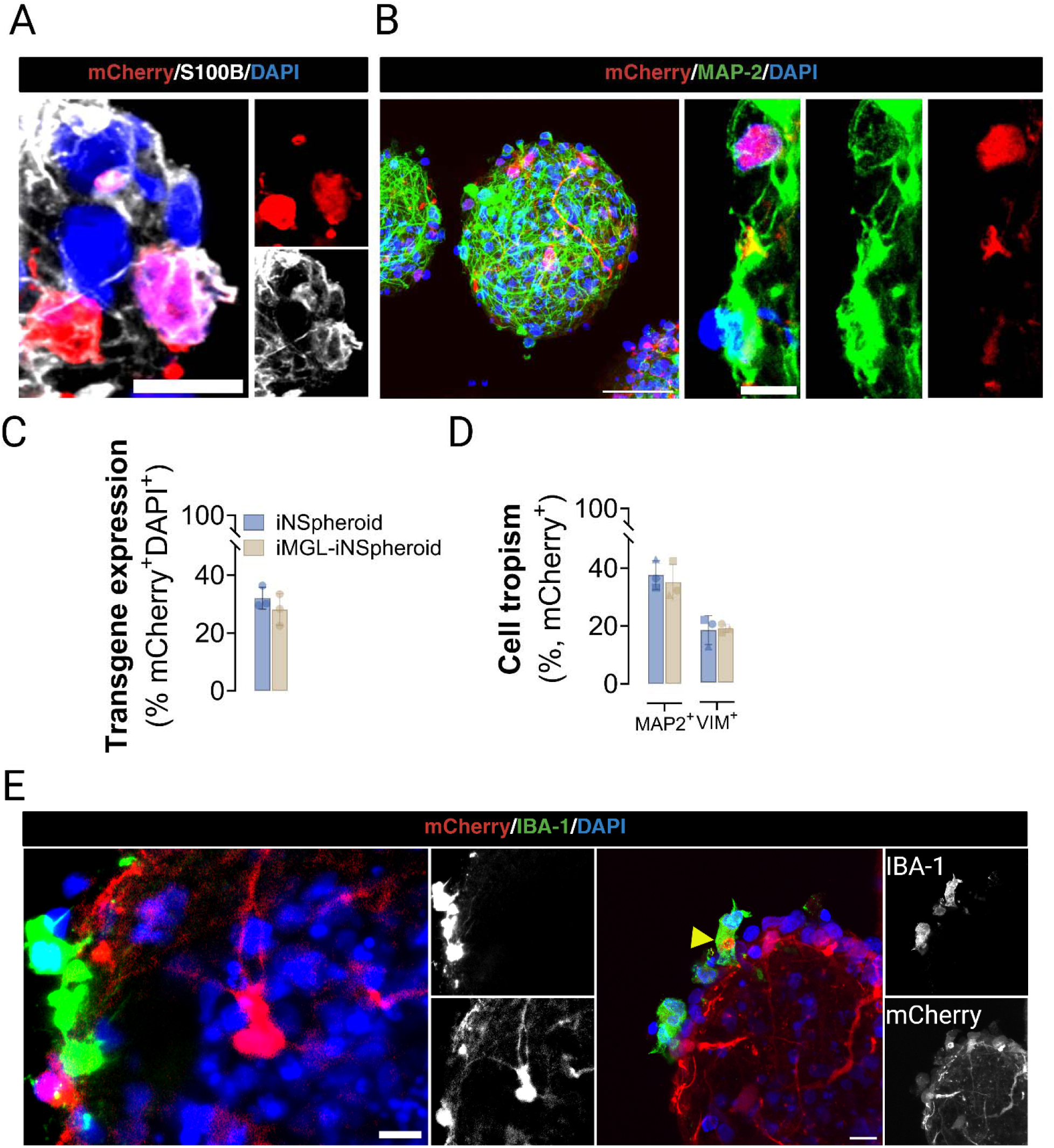
**Transduction of iMGL-iNSpheroids with rAAV9-mCherry leads to transgene expression by neural cells but not microglia.** (a) Immunofluorescence microscopy detection of mCherry (red, rAAV9 transgene) and S100B (grey, astrocytes) or (b) MAP-2 (green, neurons), for rAAV9-mCherry transduction efficiency and tropism assessment. Scale bars: 50 μm; 5 μm for zoom-in images. Immunofluorescence-based quantification of (c) mCherry-positive and DAPI-positive and (d) VIM-positive mCherry-positive or MAP2-positive mCherry-positive cells in iNSpheroids without or with iMGL (iMGL-iNSpheroids) infected with AAV9-mCherry at 5x105 VG/cell. Results are expressed as mean ± S.D., N = 3. (e) Immunofluorescence microscopy of mCherry (red, rAAV9 transgene) and IBA-1 (green, microglia). Microglia does not express the transgene; however, it clusters near mCherry-positive cells and engulfs mCherry-positive fragments (yellow arrow) .

Seven days after transduction with the rAAV9-mCherry construct, no significant difference in the percentage of cells expressing the transgene was found between conditions (iNSpheroids: 31.3 ± 3.8%, and iMGL-iNSpheroid: 29.2 ± 7.0%) (Fig. 6 C). This finding was consistent across other rAAV9 constructs carrying GFP as the transgene (Fig. S6 B). A tendency for a higher tropism of the neuronal population (MAP-2-positive cells)(iNSpheroids: 42.57 ± 3.70% and iMGL-iNSpheroids: 34.83 ± 4.24%) in comparison to the astrocytic population (VIM-positive cells) (iNSpheroids: 18.80 ± 3.10% and iMGL-iNSpheroids: 18.45 ± 3.26%) was found with all vectors under analysis, with no significant differences in the presence or absence of iMGL (Fig. 6 D and Fig. S6 A-E).

Interestingly, the mCherry transgene was not detected in any cell type at 6- and 24-hours post-transduction, by snRNA-seq, suggesting a slow progression of the virus from the engagement with the cell membrane to the cell nuclei. iMGL were not transduced with any of the rAAV9 vectors, either as 2D monocultures or in co-culture with the iNSpheroids (Fig. 6 E and Fig. S6 F and G, respectively). Nonetheless, iMGL acquired different morphological features and tended to cluster near transduced cell areas (Fig. 6 E).

Additionally, some iMGL presented phagosome-like structures (Fig. 6 E and Fig. S7 A, absence of IBA-1 signal in the cytoplasm) containing fragments of mCherry (Fig. 6 E, yellow arrowhead), which may correspond to phagocytic events. The inefficiency of rAAV to transduce immune cells, in particular microglia, has been suggested as a result of opsonization, allowing the host cell to enhance phagocytosis efficiency, blocking pathogen entry and mediating its degradation (*47*). To address the capacity of rAAV9-mCherry to enter iMGL, we evaluated the presence of AAV viral genome within iMGL by in situ hybridization using the SABER-FISH technology (*48*, *49*). By confocal microscopy, we visualized AAV genomes within iMGL cell cytoplasm and in the perinuclear region, after 24 hours of transduction, suggesting that although several rAAV9 particles are able to enter these cells, nuclear import and subsequent transduction is hampered (Fig. S7 B, C). To compare the lack of iMGL transduction with in vivo data, we injected eight-month-old C57/Bl6 wild-type mice via intracerebroventricular injection with rAAV9-mCherry (1 x 1011 VG/mouse) (Fig. S7 D). Although the overall transduction efficiency was low (13.77 ± 9.90%) rAAV9-mCherry successfully transduced 38.50 ± 18.68% of TMEM119-positive microglia, 68.99 ± 10.68% of GFAP-positive astrocytes, and 34.74 ± 2.25% and MAP2positive neurons (Fig. S7 E-G). Similarly to iMGL, CD86-positive microglia in the mouse brain were frequently found in juxta-position with rAAV9-transduced cells (Fig. S7 H) and acquired a range of different morphologies ranging from ramified to elongated forms (Fig. S7 G, I). Together, these results demonstrate that rAAV9 can transduce neurons and astrocytes in both human and mouse models, as demonstrated previously (reviewed in (*50*)). However, the discrepancy in microglial transduction between in vitro and in vivo systems suggests that species-specific differences or environmental cues may modulate rAAV9 susceptibility in microglia. This highlights an opportunity to further investigate the mechanisms regulating AAV-mediated gene delivery in human microglia, which remains a critical consideration for CNS-targeted gene therapies.

### rAAV9-mediated iMGL-iNSpheroid transduction induces a transient proinflammatory signalling

To shed light on rAAV9-host interactions in a human context, we employed our human immunocompetent iMGL-iNSpheroid model to study a critical hurdle of rAAV for clinical use: tissue innate immune response.

iMGL-iNSpheroids either untransduced (control) or transduced with rAAV9-mCherry, and collected at 0-, 6-, and 24-hours post-transduction. Given previous reports showing that excessive GFP expression can trigger vigorous inflammation and intense immune response after transduction (*51*), we focused the transcriptome analysis on the rAAV9-mCherry vector. The samples were then processed, annotated, and analysed for transcriptional changes using snRNA-seq (Fig. 7 A and Fig. S8 A-C). AAV vectors have been demonstrated to trigger innate immune pattern recognition receptors, including TLR2 and TLR9, leading to the production of inflammatory cytokines and type I interferons (IFN) (*52*). Cells within the iMGL-iNSpheroids were found to express both TLR2 and 9, demonstrating that they have the machinery to recognize these viral particles (Fig. 7 A).

**Fig. 7.**
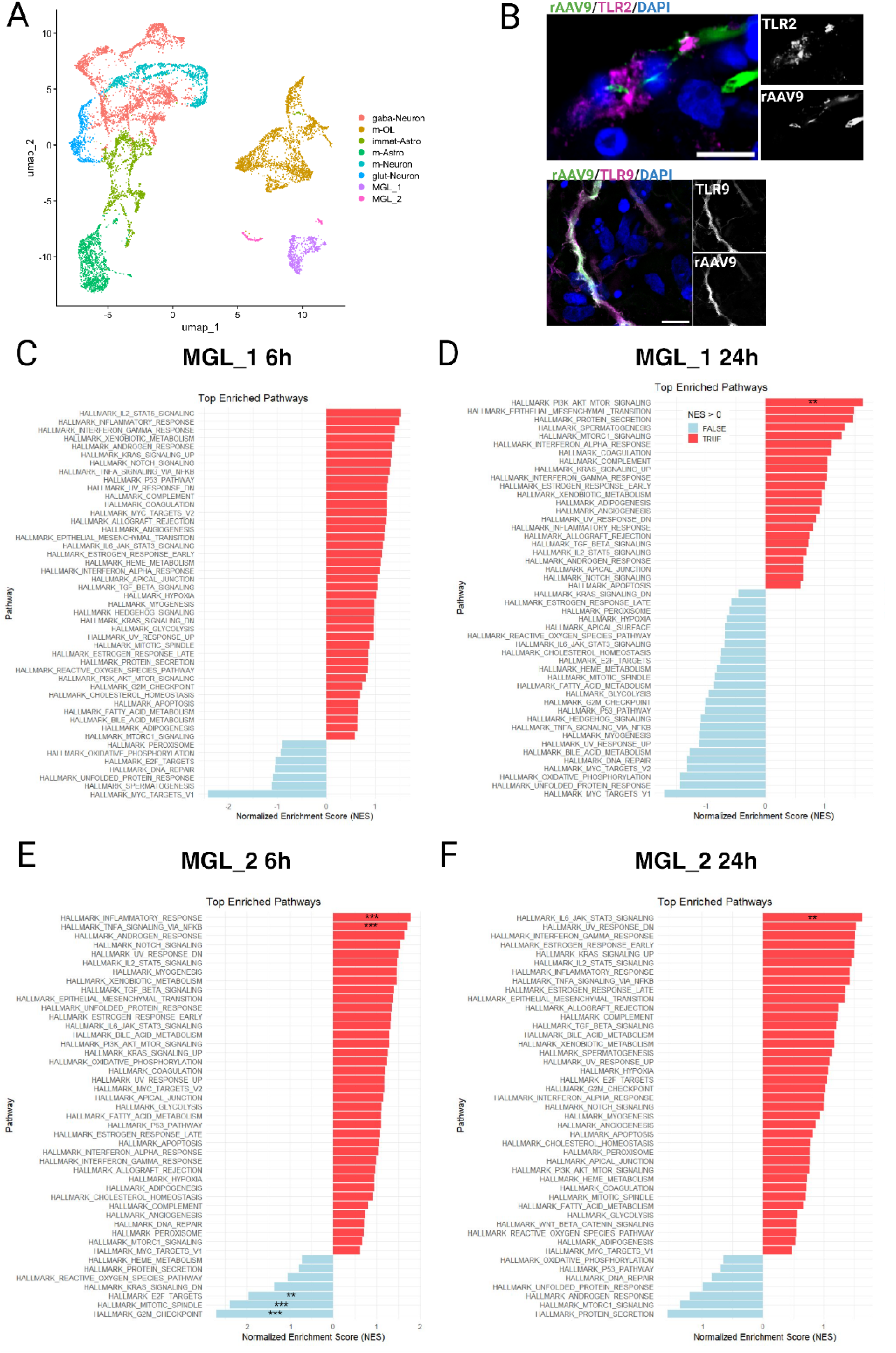
**Single-nuclei RNAseq of co-cultures (iMGL-iNSpheroids) transduced with rAAV9-mCherry reveals two different iMGL subtypes with time-dependent activation of inflammatory pathways.** (a) Clustering the integrated datasets at resolution 0.1 allowed identifying seven major clusters, corresponding to the major neural cell types (neurons, astrocytes, and oligodendrocytes) and microglia. UMAP projection, calculated with the first 30 PCs. (b) iMGL-iNSpheroids, immunostaining of toll-like receptor (TLR)2 (upper panel, magenta) and TLR9 (lower panel, magenta), and transduced cells (mCherry-positive, rAAV9, green). Nuclei are counterstained with DAPI (blue). Scale bar = 5 μm. (c) Bar plot visualizing the enriched GSEA terms in iMGL-iNSpheroid cultures transduced with rAAV9-mCherry, over non-transduced controls, against the Hallmark gene set (Molecular Signatures Database, human MSigDB - GSEA). Samples were collected at two different timepoints: 6- and 24-hours post-transduction. Top enriched pathways in sub-cluster (c) MGL (cluster 6) 6 hours post-transduction (hpt), (d) MGL (cluster 6) 24hpt, (e) MGL (cluster 7) 6hpt, and (f) MGL (cluster 7) 24hpt are illustrated. GSEA was performed on Lo g2FC pre-ranked gene lists obtained from gene expression levels of the different cell types. NES, normalized enrichment score; *, adjusted p< 0.05; **, adjusted p< 0.01; ***, adjusted p< 0.001.

Interestingly, two different iMGL subclusters were identified. Gene set enrichment analysis revealed that the two annotated iMGL clusters (Fig. 7 B and Fig. S8 A-C, clusters 6 and 7) exhibited distinct responses at both 6- and 24-hours post-transduction. Cluster “MGL_1” showed a significant upregulation of the “PIK3/Akt/mTOR pathway” at 24 hours, along with a pronounced downregulation of “MYC targets” at both time points (Fig. 7 C, D). In contrast, cluster “MGL_2” exhibited a strong inflammatory response at 6 hours post-transduction, including a significant upregulation of hallmark gene sets such as “inflammatory response” and “TNF via activation of NF-κB”, as well as a downregulation of pathways involved in DNA damage repair, such as “G2M checkpoint”. Interestingly, this inflammatory response was no longer observed at 24 hours post-transduction. Instead, an upregulation of the “IL-6/JAK/STAT3 pathway” emerged, which can be a downstream product of the activation of “TNF via activation of NF-κB” (Fig. 7 E, F).

These distinct response profiles may be linked to microglial plasticity in response to inflammatory stimuli. Indeed, we observed MGLs with different morphological characteristics, which may correspond to distinct transcriptomic profiles. When analyzing target genes associated with key microglial functions, such as phagocytosis (CD68, TREM2) and inflammatory responses (C3, TNF, CXCL8), we found a trend toward increased expression of inflammatory genes in cluster “MGL_1”, whereas cluster “MGL_2” exhibited higher expression of phagocytosis-related genes (Fig. S8 D, E).

The macroglial cells within the iMGL-iNSpheroids, including astrocytes (Fig. S9) and oligodendrocytes (Fig. S10), exhibited a significant upregulation of the “TNF via activation of NF-κB” pathway at 6 hours post-transduction. Interestingly, by 24 hours, astrocytes displayed increased expression of genes associated with epithelial-to-mesenchymal transition (EMT) and myogenesis pathways, a phenomenon previously observed in astrocytes activated in response to glioma (*53–58*). Neurons did not exhibit activation of inflammatory response pathways. However, at 6 hours post-transduction, the ’mature neuron’ cluster showed significant downregulation of MYC targets, while the ’GABAergic neuron’ cluster exhibited reduced expression of DNA repair pathways (Fig. S11).

To validate the snRNA-seq findings, a pilot bulk proteomics and targeted secretome analyses were performed (Fig. 8 and Fig. S12). Whole proteome analysis of iMGL-iNSpheroids, comparing rAAV9-transduced versus control samples at 48 hours post-transduction, identified several proteins involved in viral infection and immune system activation (Fig. S12 A, B). Correlation analysis between the pseudobulk snRNA-seq dataset and bulk proteomics revealed significant associations between multiple differentially modulated genes and proteins (Fig. S12 C). Notably, genes linked to the innate immune response to viral infection (e.g., TLR2, IRAK3, TLR7, SOCS1, CLU) correlated with corresponding proteins (e.g., ISG15, CLEC4C, C9), along with other immune-related targets (e.g., MYH9, RAB31, IQGAP1, MMP12, CD63) (Fig. S12 C, D, E). Additionally, secretome analysis demonstrated that rAAV9-mCherry transduction of iMGL-iNSpheroids led to the secretion of CXCL10, IFN-γ, and CCL2 (Fig. 8 A). These proteins are regulated by the IFN/STING and NF-κB pathways and are well known to be secreted in response to viral infection (*59*, *60*). In contrast, little to no differential secretion of proteins was observed in iNSpheroids alone compared to their respective untransduced controls. Interestingly, similar trends were found when comparing the different rAAV9 vectors used (Table S3). Altogether, these findings highlight the critical role of microglia in CNS immune defense, as they actively recognize, respond to, and eliminate external entities such as rAAV (*61–66*).

**Fig. 8.**
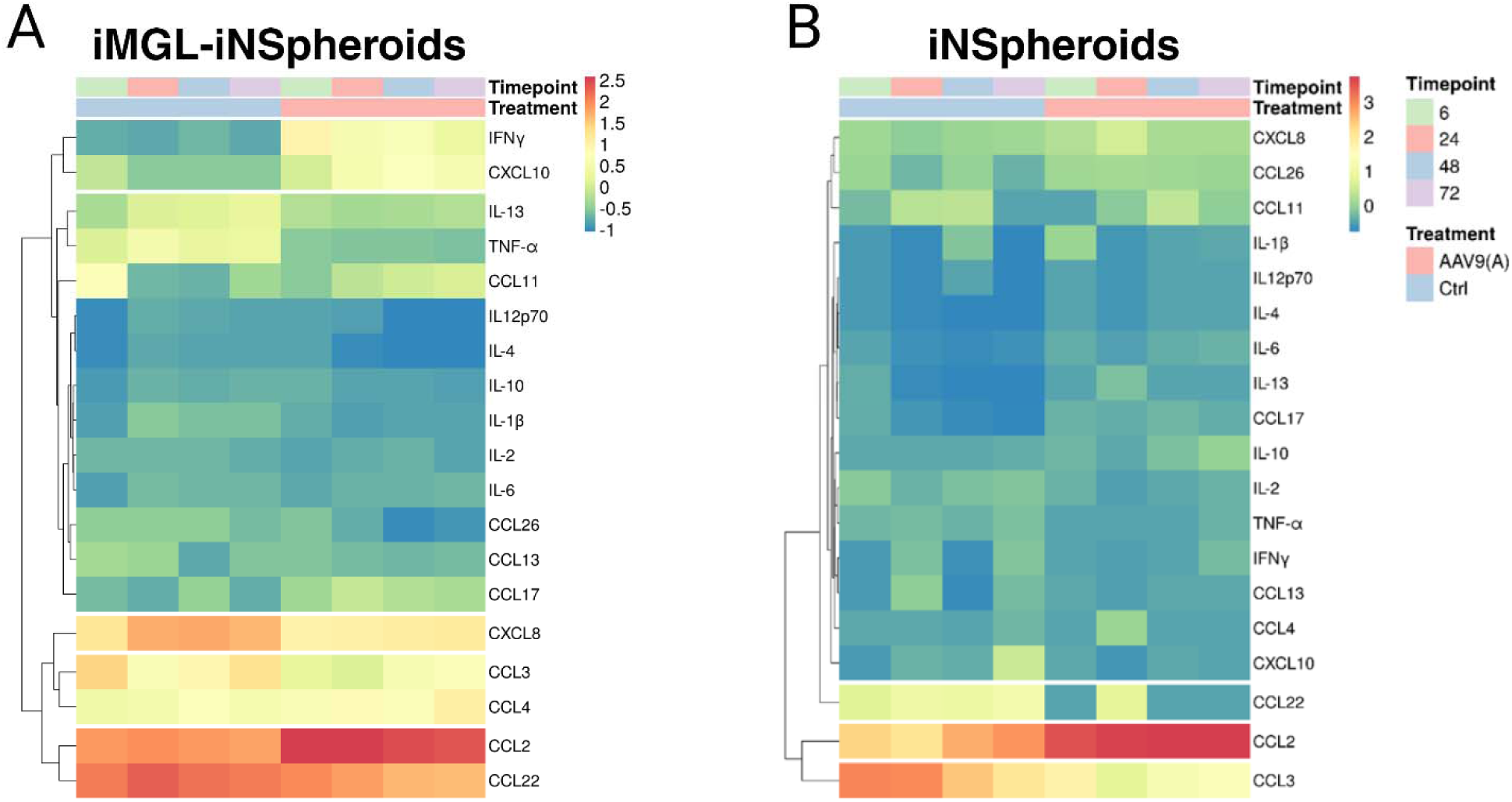
**iMGL-iNSpheroids secrete proinflammatory cytokines and chemokines upon rAAV9-mCherry transduction.** (a) Hierarchical clustering of the protein targets identified by secretome analysis of the iMGL-iNSpheroids and (b) iNSpheroids. Data represented as mean of N = 4 (iPSC(IMR90)-clone 4).

## Discussion

This study describes the development and application of a novel 3D hiPSC-derived innate immunocompetent CNS model, the iMGL-iNSpheroids, for the preclinical assessment of rAAV-based gene therapies. This model relies on the computer-controlled bioreactor co-culture of hiPSC-derived microglia with iNSpheroids composed of neurons, astrocytes, and oligodendrocytes, thereby providing a comprehensive platform to study the neuro-immune axis in a physiologically relevant context. Our findings underscore the critical role of microglia in modulating neuroinflammatory responses and highlight the potential of this model to enhance our understanding of rAAV9 immunogenicity within the CNS. The ability to sustain a constant composition of homeostatic microglia and replicate complex cellular interactions and innate immune responses of the human CNS in vitro iNSpheroids, represents a significant advancement in the field, offering a more accurate and predictive tool for preclinical research.

The iMGL-iNSpheroid model successfully sustained integration and functional interaction of microglia with other neural cell types, as evidenced by the expression of microglial markers such as IBA-1, P2RY12, and TMEM119. The presence of microglia within the iNSpheroids was confirmed through immunolabelling and transcriptomics analyses, which revealed the expression of genes associated with phagocytosis, lysosomal function, and lipid metabolism. These findings suggest that the microglia within the iMGL-iNSpheroids retain their functional characteristics and can effectively engage in synaptic pruning and debris clearance, akin to their in vivo counterparts. The ability of microglia to dynamically respond to environmental cues and maintain homeostasis within the CNS underscores their importance in neuroinflammatory processes and highlights the relevance of including microglia in in vitro models to study CNS pathophysiology.

rAAV as a gene therapy vehicle has demonstrated potential in treating CNS disorders (*67*). The initial promising results from intravenous rAAV-based gene therapies targeted the liver for various metabolic diseases (*68*). Using the same delivery strategy to target the adult CNS has several advantages, namely the decreased invasiveness (*69*). However, the high dosages needed for this administration route can lead to immune responses and toxicity. Recently, Au and colleagues have reported that amongst 255 rAAV-based gene therapy clinical studies, the most prominent serious adverse events were hepatotoxicity, thrombotic microangiopathy and neurotoxicity (*70*). Indeed, neuroinflammation as a result of vector transduction can significantly reduce therapeutic efficacy (*13*, *71*). Most preclinical trials have relied on animal models, which, due to the inherent differences in their immune systems, resulted in limited translatability to humans (*13*, *71*). However, more clarification is needed to understand and potentially avoid significant anti-rAAV immune response and adverse events. Most preclinical trials have relied on animal models, which, due to the inherent differences in their immune systems, resulted in limited translatability to humans 12,69. However, developing a human in vitro model to study microglia has been challenging with most protocols to date reporting microglia activation or limited capacity to retain them for long periods (reviewed in (*72*, *73*)). Human iPSCs have shown to be a valuable methodology enabling the generation of microglia from both healthy donors and patients (*15*, *17*, *74*) (reviewed in (*75*)).

rAAV serotype 9 effectively transduced the iNSpheroids, with and without iMGL, with tropism for both neuronal and astrocytic populations. This tropism was further validated in a C57/Bl6 wild-type mouse model, where a high percentage of the GFAP-positive astrocytes were transduced. Interestingly, iMGL within the iNSpheroids model were not transduced by rAAV9, in line with previous reports suggesting that immune cells are less susceptible to AAV transduction, likely due to opsonization and subsequent phagocytosis of AAVs (reviewed in (*76*, *77*)). In contrast, TMEM119-positive microglia were transduced in the mouse brains, suggesting potential species-specific differences or environmental factors influencing transduction efficiency. In the iMGL-iNSpheroids model, microglia were frequently found in close proximity to transduced cells, suggesting their involvement in the innate immune response elicited by the rAAV9 tested.

The transcriptional and proteomics analyses of iMGL-iNSpheroids transduced with rAAV9 revealed the activation of several inflammatory pathways, including IFN-α and −γ responses, TNF-via activation of NF-κB, STING through p53 activation. Two different cell clusters were annotated as microglia and were found to have different expression profiles for genes associated with phagocytosis and innate immune response. Astrocytes and oligodendrocytes also assembled a response against rAAV9 transduction, activating KRAS signalling after 6 hours, which has been previously linked with immune signalling in several types of cancer (reviewed in (*78*)). Proteomics analysis corroborated this activation profile with the expression of several proteins involved in the innate immune response against viral infection, namely C9, ISG15, and in tissue injury and remodelling (e.g., MMP12). These findings align with the known mechanisms of innate immune recognition of viral vectors and underscore the relevance of the iMGL-iNSpheroid model in studying the molecular underpinnings of rAAV9 immunogenicity (*61–66*). The transient nature of the inflammatory response, with mild differences in the response between 6- and 24-hours post-transduction, suggests a dynamic interplay between the different cellular counterparts. These results support previous findings showing that innate immune responses to rAAV are mild and transient (reviewed in (*79–81*)).

The cGAS-STING axis is also increasingly implicated in the cellular response to DNA damage (reviewed in (*82*, *83*)). However, its activation during AAV-mediated transduction in the CNS remains poorly characterised. Current evidence indicates that both wild-type and recombinant AAV vectors can induce DNA damage responses, as demonstrated by their colocalization with DNA damage foci in nuclei of HeLa and MRC5 cells (*84*) and the activation of p53-mediated signalling in human haematopoietic stem and progenitor cells (*85*). In iMGL-iNSpheroids, we observed the modulation of several pathways involved in DNA damage and repair across most cellular components, including MYC targets, the G2M checkpoint, and E2F targets. These results suggest that iMGL-iNSpheroids quickly sense and respond to rAAV9 upon contact with viral particles, leading to detectable cellular stress and damage responses as early as 6 hours post-exposure.

We also explored the impact of microglia on neuronal maturation and synaptic activity within the iNSpheroids. The presence of microglia was associated with enhanced synaptic vesicle trafficking, as indicated by the increased fluorescence intensity and faster decay of FM 1-43 dye in iMGL-iNSpheroids compared to iNSpheroids alone. This observation aligns with the known role of microglia in regulating synaptic function (reviewed in (*86*)) and underscores their contribution to the maturation of neural circuits within the 3D model. The co-culture with iMGL significantly enhanced synaptic vesicle trafficking compared to iNSpheroids alone, indicating improved neuronal functionality. The interactions between microglia and synapses, evidenced by the co-localization of IBA-1 and PSD-95 and GEPH signals, suggest active microglial engagement in regulating synaptic morphology by eliminating excess or dysfunctional synapses. This finding aligns with the known role of microglia in preventing neuronal hyperexcitability and maintaining homeostasis within the CNS (*87*). The ability of microglia to modulate synaptic activity and promote neuronal maturation highlights their critical role in the development and maintenance of functional neural networks, which is essential for the accurate modelling of CNS diseases and therapeutic interventions (reviewed in (*86*, *88*, *89*)).

Microglia play a pivotal role in maintaining CNS homeostasis by constantly surveying the brain parenchyma to identify, eliminate, and resolve inflammation. Upon pathogen recognition, microglia initiate and sustain neuroinflammation, recruiting additional microglia and astrocytes to the injury site (reviewed in (*90*)). Microglia within the iNSpheroids exhibited a variety of morphologies, ranging from ramified to more rounded forms with an increased cell soma and lower branch density. The ramified morphology is characteristic of non-activated resting microglia, primarily surveilling and maintaining tissue integrity. In contrast, the amoeboid shape reflects increased motility and proliferative capacity, typically observed in microglia responding to the presence of cellular damage and/or proinflammatory cues (*34*, *91–93*). Alongside their phagocytic capacity and involvement in synaptic pruning, this morphological plasticity underscores the functional adaptability of microglia within the brain (*88*, *94*). Interestingly, we found amoeboid microglia cells close to dead cells and regions where rAAV9-transduced cells were localized, demonstrating microglial morphofunctional plasticity when encountering cellular damage. Given that neurons release chemotactic signals to recruit microglia for the clearance of apoptotic cells and defective synapses, the observation of rounded IBA-1-positive cells near neurons, and the expression of late-endosomes/phagocytosis-related genes and proteins (e.g., CD68, TREM-2, TYROBP and CLEC7A), suggests active microglial engagement in phagocytic activity, which is essential for the formation of functional and mature neural circuits (*95–97*).

Timely removal of dying cells, debris or pathogens by phagocytes, like microglia, is essential to maintaining host homeostasis. Phagocytosis is a process partially regulated by the balance of “eat me” and “don’t eat me” signals expressed on the surface of cells.

Upon contact, eat me signals activate “pro phagocytic” receptors expressed on the phagocyte membrane and signal to promote phagocytosis. Conversely, “don’t eat me” signals engage anti phagocytic receptors (e.g., SIRPα/ CD47) to suppress phagocytosis. The complement cascade has been for long known as a crucial pathway in regulating synaptic pruning, governing the fine balance between the “eat-me” and “don’t-eat-me” signalling between microglia and neurons (*98–100*). In our study, we demonstrate that some neurons express C3 and C1QA, localized on the cell membrane (cellular damage/death) or in puncta, together with the concomitant expression of C1QA-C and C3 at the transcript level in glutamatergic neurons and in microglia.

The ability to study the dynamic interactions between microglia and neural cell types in response to inflammatory stimuli offers valuable insights into the mechanisms of disease and the potential impact of therapeutic interventions on these processes. Upon exposure to proinflammatory stimuli, such as TIC and LPS plus IFN-γ, the iMGL-iNSpheroids exhibited a robust neuroinflammatory response, characterised by the upregulation of several immunomodulators at both gene and protein secretion levels, including IL-6, CXCL8, and CCL2. These stimuli are known to be produced during pathogen infection and neurological disorders such as Alzheimer’s disease (*34*, *101*). Notably, the presence of microglia modulated the inflammatory response, as evidenced by the differential gene expression and protein secretion profiles in iMGL-iNSpheroids compared to iNSpheroids alone, especially under LPS plus IFN-γ stimuli. This finding highlights the importance of microglia in shaping the neuroinflammatory landscape and suggests that their inclusion in the model provides a new avenue to study neuroinflammatory processes, more accurately mimicking the human CNS neuro-immune response.

Overall, our work presents a novel methodology to incorporate microglia, maintain a homeostatic phenotype, communicate with cells from the microenvironment, and improve neuronal functionality. The iMGL-iNSpheroids model demonstrated its suitability as complementary tools for rAAV-based gene therapy preclinical research by offering a versatile and simpler platform to uncover innate immunity mechanisms upon rAAV challenge. Considering the limitations of the current in vitro and in vivo models in recapitulating the human CNS innate immunity, we envision this model as offering a simplified and robust vision of the human neuro-immune context, allowing for higher throughput in preclinical rAAV screening. While not averting the clinical utility of rAAV vectors for the treatment of CNS disorders, this work highlights the relevance of cell-intrinsic innate immune activation as a potential mediator of AAV immunogenicity and, if required, may inform the future immunomodulatory strategies improving outcomes of CNS-targeted AAV-based gene therapies.

## Materials and Methods

### Human induced pluripotent stem cell (hiPSC) culture

Two commercially available hiPSC lines were used in this work, iPSC(IMR90) clone 4 and iPSC(DF6-9-9T.B) (WiCell). The hiPSC were expanded on Matrigel hESC-Qualified Matrix, LDEV-free (Corning, cat. no 354277), in mTeSR™1 cGMP medium (StemCell Technologies, cat. no 85850) under feeder-free culture conditions. A complete medium exchange was performed every day. Cells were maintained in a humid atmosphere with 5% CO_2_, at 37 °C.

### Mouse models and in vivo experiments

All animal experiments were conducted in accordance with the European Community Council Directive (2010/63/EU) for the care and use of laboratory animals and previously approved by the Responsible Organization for the Animals Welfare of the Faculty of Medicine and Center for Neuroscience and Cell Biology of the University of Coimbra (ORBEA and FMUC/CNC, Coimbra, Portugal, ORBEA_66_2015/22062015 and ORBEA_289_2021/10122021). Researchers performing animal experiments received appropriate training (FELASA-certified course) and certification from Portuguese authorities (Direcção Geral de Alimentação e Veterinária).

C57BL/6 wild-type mice were housed in a temperature-controlled facility on a 12h light/12h dark cycle with water and food provided ad libitum. For stereotaxic injection, mice were anaesthetised with 2 % isoflurane (IsoVet 1000 mg/g) and maintained under 1.2 % isoflurane with 0.8 L/min oxygen using the RWD anaesthesia system. AAV9-mCherry was injected into the right lateral ventricle using the following stereotaxic coordinates relative to bregma: anteroposterior 0.3 mm, lateral: 1 mm, ventral: 2,5 mm and tooth bar 0. A 25 mL hamilton syringe with a Small Hub RN Needle (point style 2) was used to inject a total volume of 8 mL at an infusion rate of 2 mL/min. After completing the infusion, the syringe was withdrawn 0.3 mm and kept in place for an additional 3 minutes. Following the procedure, animals were sutured using a 4/0 Silkam suture (B. Braun) and administered buprenorphine for analgesia.

Mice were euthanized for four weeks post-injection. Terminal anesthesia was induced using a mixture of ketamine (Clorketam 1000, Vétoquinol) and xylazine (Rompun, Bayer), followed by transcardial perfusion with ice-cold PBS (1×, pH 7.4). The left hemisphere was post-fixed for immunofluorescence in 4% paraformaldehyde (PFA) in PBS for 24 hours at room temperature, then cryoprotected in 25% sucrose for 48 hours before storage at –80°C.

### Generation of NPCs from hiPSC

iPSC(IMR90)-clone 4 and iPSC(DF6-9-9T.B)cells were plated at 0.8 x 10^5^ cell/cm^2^ in mTeSR™1 cGMP medium and allowed to adhere to Matrigel hESC-Qualified Matrix-coated plates for 24 hours. Afterward, cells were cultured in either (i) DualSMADi culture medium (iPSC(IMR90)-clone 4): N2B27 medium supplemented with SB431542 at 1 mM and LDN193184 at 10 mM (both from STEMCELL Technologies); or (ii) STEMdiff™ Neural Progenitor Medium (STEMCELL Technologies, cat. no. 05833) (iPSC DF6 9-9T.B). N2B27 medium was composed by DMEM/F-12 and Neurobasal in 1:1 ratio, N2 supplement (1x), B27 supplement without vitamin A (0.5x), GlutaMAX (1x), Penicillin-Streptomycin (1%), MEM Non-Essential Amino Acids Solution (1%) and 2-mercaptoethanol (50 μM) (all from Gibco™), and Insulin (20 μg/mL) (Sigma-Aldrich). A 100% medium exchange was performed daily, for 7 (DF6-9-9T.B) or 10 (iPSC(IMR90)-clone 4) days. On day 7 or 10, cells were recovered and plated on poly-L-ornithine-laminin (PLOL)-coated surfaces. PLOL coating was prepared by performing a 3-hour incubation at 37 °C with 0.16 mg/mL poly-L-ornithine in PBS (with Ca^2+^ and Mg^2+^), followed by a washing step and a 3-hour incubation at 37 °C with 1 μg/mL laminin in PBS (with Ca^2+^ and Mg^2+^). Cells were maintained in DMEM/F12 medium with Glutamax (Life Technologies) supplemented with 1% N2 supplement (Life Technologies), 0.1% B27 supplement (Life Technologies), 1.6 μg/mL glucose (Sigma-Aldrich), 20 μg/mL insulin (Sigma-Aldrich), 20 ng/mL rhu-bFGF (Peprotech)(*23*). The iPSC(IMR90) clone 4 line was maintained on PLOL-coated surfaces in an NPC expansion medium (EM), composed of DMEM/F12 medium with Glutamax (Life Technologies) supplemented with 1% N2 supplement (Life Technologies), 0.1% B27 supplement (Life Technologies), 1.6 μg/mL glucose (Sigma-Aldrich), 20 μg/mL insulin (Sigma-Aldrich), 20 ng/mL rhu-bFGF (Peprotech) and 20 ng/mL rhu-EGF (Sigma-Aldrich). The medium was changed every other day until confluency. NPCs derived from the two hiPSC lines were passaged at 90-100% confluence (typically every 3-4 days). Cells were dislodged by 0.05% Trypsin-EDTA (1-2 min), which was neutralized with DMEM supplemented with 10% FBS (Life Technologies). Cells were sedimented by centrifugation, resuspended in EM, and plated on PLOL-coated T-flasks, at a cell density of 3 x 10^4^ cell/cm^2^. A 50% medium exchange was performed on day 2 of culture. Cell concentration and viability were determined by the trypan blue exclusion method, in a Fuchs-Rusenthal haemocytometer. Cells were maintained under a humidified atmosphere, in a multi-gas cell incubator (Sanyo), with 5% CO_2_ and 3% O_2_, at 37 °C.

### Erythromyeloid progenitor differentiation from hiPSC (iEMP)

hiPSC lines were differentiated into EMPs using a STEMdiff™ Hematopoietic Kit (STEMCELL Technologies, cat. no. 05310). One day before induction of differentiation, hiPSCs were harvested through incubation with ReLeSR™ (STEMCELL Technologies cat. no. 05872) according to the manufacturer’s instructions to obtain small colonies (100– 200 μm in diameter). Then, the colonies were seeded at 20 aggregate/cm^2^ in mTeSR™1 supplemented with 10 μM Y-27632. After one day, the mTeSR™1 medium was replaced with Medium A (STEMdiff™ Hematopoietic Basal Medium containing 0.5% (v/v) of Supplement A) to induce mesoderm differentiation (day 0 of differentiation). On day 2 of differentiation, half of the medium was changed to fresh Medium A. On the following day, the medium was completely changed to Medium B (STEMdiff™ Hematopoietic Basal Medium containing 0.5% (v/v) of Supplement B). Half medium exchanges were performed on days 5, 7, 8, and 10 to promote further lineage commitment towards erythromyeloid cells. By day 12, iEMPs could be harvested from the culture supernatant and froze at 1-2 x10^6^ cell/ mL in BamBanker (LYMPHOTEC Inc., cat. no. 302-14681).

### Microglia differentiation from iEMP (iMGL)

On day 0 of iPSC-derived microglia (iMGL) differentiation, iEMP were plated at 1-2 x 10^4^ cell/cm^2^ on Matrigel hESC-Qualified Matrix (1 mg/mL). iMGL differentiation was induced by employing the STEMdiff™ Microglia Differentiation and Maturation kits (STEMCELL Technologies, cat. no. 100-0019 and 100-0020, respectively), following manufacturers’ instructions. Throughout the differentiation of iEMP to iMGL, cells predominantly grow non-adherently. On days 2, 4, 6, 8, and 10, half of the volume was replenished by slowly removing the spent medium and adding fresh one. On day 12, suspension cells were collected and centrifuged at 400 xg for 5 min, at room temperature (RT). Cells were then resuspended in a 1:1 mixture of conditioned and fresh medium.

After that, fresh medium (half the initial volume) was supplemented every other day until day 24. On day 25, cells were centrifuged as previously and resuspended in the maturation medium, following the same feeding regime. On day 28, cells were collected for iMGL phenotype characterization and processed for co-culture experiments.

### hiPSC-derived neurospheroid (iNSpheroid) differentiation

This protocol follows our previously published methodology in (*23*, *24*). hiPSC-NPCs were expanded and harvested as described in the previous section. The resulting cell suspension was passed through a 70 μm nylon strainer (Millipore) before bioreactor inoculation, to eliminate cell clumps. The obtained single-cell suspension was diluted to a cell density of 4 x 10^5^ cell/mL in aggregation medium (AM), which is composed of EM with reduced EGF/FGF concentration (5 ng/mL) and supplementation of 5 μM Y-27632. The single cell suspension was inoculated into a software-controlled stirred-tank DASGIP Bioblock bioreactor system (Eppendorf), as described previously(*23–26*). Culture conditions were set to maintain cells under 3% dissolved oxygen (15% of air with 21% oxygen), pH 7.4, 37 °C, and a stirring rate of 70 rpm. To control the aggregate size and avoid aggregate fusion, the stirring rate was gradually increased up to 90 rpm, with 10 rpm steps, based on visual inspection of the culture. After 72 hours of culture, the perfusion operation mode was activated, with a dilution rate of 0.33 day^-1^ (i.e., 33% working volume exchange per day), under gravimetric control. To prevent the loss of aggregates through the outlet perfusion line, a metallic filter of 20 μm pore size was adopted as a cell retention device(*25*). After a 7-day aggregation period, differentiation was induced by replacing the AM perfusion medium with differentiation medium (DM) and maintaining the culture for an additional 23 days (a total of 30 days). DM was prepared by supplementing DMEM/F12 with Glutamax with 2% B27 supplement, 1.6 μg/mL glucose, 10 μg/mL insulin, 10 μg/mL putrescin, 63 ng/mL progesterone, 50 μg/mL apotransferrin, 50 ng/mL sodium selenium (all from Sigma-Aldrich) and 200 mM ascorbic acid (Wako).

### Ambr®15 Cell Culture Bioreactor System

Prior to inoculation, the vessels of the Ambr^®^ 15 Cell Culture Bioreactor (from here onwards referred to as microbioreactor) were loaded onto the platform, and stabilized for pH (7.4), temperature (37 °C) and pO_2_ (15%), as described previously(*26*). The working volume range was 11-15 mL, and the impeller speed was set to 300 rpm, in down-pumping mode. The iNSpheroids were collected from the software-controlled stirred-tank DASGIP Bioblock bioreactor system (Eppendorf), sedimented and resuspended in fresh medium prior to the microbioreactor inoculum. iNSpheroids were inoculated at 1.0 x 10^6^ cell/mL. An iMGL suspension at 0.1 x 10^6^ cell/mL was added subsequently to the co-culture vessels. Cells were co-cultured for three days prior to downstream analysis or inflammatory challenge. The culture medium was composed of a 1:1 mixture of DMEM/F-12 without phenol red with 0.1X Glutamax and Neurobasal™ Medium (Gibco™, cat. no. 21103049), 1% B27 supplement, 1.6 μg/mL glucose, 10 μg/mL insulin, 10 μg/mL putrescin, 63 ng/mL progesterone, 50 μg/mL apotransferrin, 50 ng/mL sodium selenium (all from Sigma-Aldrich), 200 mM ascorbic acid (Wako), recombinant human M-CSF, IL-34 and TGF-JJ (all 10 ng/mL, Peprotech, 300-25, 200-34, 100-21, respectively).

### Cell concentration and viability

Cell concentration and viability were determined using NucleoCounter NC-200, an automated imaging counter, according to the manufacturer’s instructions.

### Cell viability, aggregate concentration, and size determination

Cell viability was assessed using fluorescein diacetate (FDA) and propidium iodide (PI) dual staining. The aggregates were incubated with 20 μg/mL FDA (Sigma-Aldrich, cat. no. F7378) and 10 μg/mL PI (Sigma-Aldrich, cat. no. P4864) in Dulbecco’s phosphate-buffered saline without Ca ^2+^ and Mg^2+^ (DPBS (-/-), Gibco™, cat. no. 14190169) and visualized using a fluorescence microscope (DMI6000, Leica). Images were acquired with a monochrome digital camera (Leica DFC360 FX) and processed using ImageJ software version 1.54j (https://imagej.net/ij/).

Aggregate concentration was assessed by manual counting using a phase-contrast microscope. Aggregate size was determined through the analysis of FDA fluorescence images leveraging from ImageJ. The area occupied by the aggregates was defined through manual threshold adjustment, and the Feret diameter (in μm) was measured.

### Neuroinflammation induction

Cytokine concentrations and timepoints for examining the iNSpheroids cell culture were selected based on previous studies from our laboratory and others that reported effects on glial cell responses(*34*, *102*). Based on the gene expression and protein secretion dynamics reported in the literature, we examined iNSpheroid responses from 6-72 h after (i) TNF-α (30 ng/mL, Cell signalling Technology, 8902SF), IL-1α (3 ng/mL, Sigma, I3901) and C1Q (400 ng/mL, MyBioSource, MBS143105), (ii) LPS (10 ng/mL, Lipopolysaccharides from Escherichia coli, L4516-1MG, Sigma-Aldrich) and IFN-gamma (IFN-y, 50 ng/mL, Peprotech, 300-02), and (iii) recombinant human IL-4 and IL-13 (20 ng/mL, Peprotech, 200-04 and 200-13, respectively). The different cocktails were prepared in iMGL-iNSpheroid medium, and 1 mL was added to the respective STB vessel (iNSpheroids with and without iMGL) or well plate (iMGL alone). The stimulus was maintained throughout 7 days.

### Viral vector production and purification

In this work, three different rAAV9 vectors were used. rAAV9-eGFP particles were supplied by the Bioproduction Unit of iBET. Two different plasmid cassettes were used, and vector production was attained by triple transfection in the human embryonic kidney (HEK)293T expression system and purified by affinity chromatography. Concentration and buffer exchange were performed against PBS containing CaCl_2_ and MgCl_2_ (Gibco™, cat. no. 14040091). rAAV stock titters were determined by the real-time quantitative PCR (qPCR) titration method (*103*).

rAAV9-mCherry used is part of the joint efforts of the IMI ARDAT (Accelerating Research and Development for Advanced Therapies), under the grant agreement ID 945473.

The multiplicity of infection (MOI) was determined as the number of viral genomes per cell (VG/cell) for rAAVs.

### rAAV transduction and tropism assessment

For all rAAV preparations, 5 x 10^5^ VG/cell was used based on previous reports(*104*). rAAV9 (1) and (2) encoded enhanced green fluorescent protein (eGFP) under the CMV and CAG promoters, respectively. rAAV9 (3) encoded mCherry fluorescence protein under the CAG promoter.

iMGL and iNSpheroids, with and without iMGL, were maintained in culture up to 7 days post-transduction, with medium replenishment after sampling (culture dilution).

### Immunofluorescence microscopy

iNSpheroids, with or without iMGL, were harvested after 3 days of co-culture and 7 days of transduction, washed for three times with DPBS (with Ca^2+^ and Mg^2+^), and fixed in 4% paraformaldehyde (PFA) plus 4% sucrose in PBS (+/+) for 20 min at RT. Afterwards, spheroids were washed three additional times with DPBS. Immunostaining protocol was carried out as previously described(*104*). Cell nuclei were counterstained with DAPI diluted in DPBS, for 5 min (1:1000, Life Technologies), followed by three washes in DPBS. Cells were mounted between a coverslip and a glass slide with ProLong™ Gold

Antifade Mountant (Life Technologies). Images were acquired on a Zeiss LSM880-point scanning confocal microscope controlled with the Zeiss Zen 2.3 (black edition) software. Images were processed using ImageJ software and only linear manipulations were performed.

For morphological analysis, IBA1-positive iMGL were singled-out and projected in 2D. Cell count function in the software AIVIA v14.1.0 was applied, enabling the extraction of area, length and breadth of the cell measurements.

### RT-qPCR

cDNA from 100000 cells was obtained employing Power SYBR™ Green Cells-to-CT ™ Kit (ThermoFisher, cat. no. A353380) according to the manufacturer’s instructions. qPCRs were performed in triplicates using LightCycler 480 SYBR Green I Master Kit (Roche) and the primers listed in Table S1. The reactions were performed with LightCycler 480 Instrument II 384-well block (Roche). Quantification cycle values (Cq’s) and melting curves were determined using LightCycler 480 Software version 1.5 (Roche). All data were analysed using the 2^−ΔΔCt^ method for relative gene expression analysis(*105*). Changes in gene expression were normalized using the housekeeping genes RPL22 (ribosomal protein L22), GADPH (Glyceraldehyde 3 Phosphate Dehydrogenase), and HPRT1 (Hypoxanthine Phosphoribosyltransferase 1) as internal controls. Statistical analysis was carried out using GraphPad Prism v10.1.1 software.

### MSD assay

Three pre-coated MSD (Meso Scale Discovery) plates were, namely V-PLEX Neuroinflammation Panel 1, Human Kit and V-PLEX Cytokine Panel 1 Human Kit and V-PLEX Chemokine Panel 1 Human Kit (cat. no K15210D, K15050D and K15047D, respectively). Recombinant proteins (standards) or iNSpheroid (with and without iMGL) supernatants were diluted by serial dilution in a polypropylene 96 well plate (low bind), to attain the desired concentrations using the assay diluent provided. Samples were then transferred in duplicates to each well of the antibody-coated MSD plate and incubated with shaking for 1 h at RT. After the disposal of samples and four wash cycles with 35 μL of wash buffer each, 10 μL of the primary detection antibody (SULFO-TAG (ST) labelled or non-labelled) was added to each well and incubated with shaking for 1 h at RT. For non-labelled primary detection antibodies, 10 μL goat anti-mouse SULFO-TAG labelled secondary detection antibodies (1:1 000 in blocking buffer; Meso Scale Discovery) were added to each well after an additional washing step and incubated with shaking for 1 h at RT. After washing three times with wash buffer, 35 μL of read buffer T with surfactant (Meso Scale Discovery) was added to each well and the plate was imaged on a Sector Imager 6000 (Meso Scale Discovery) according to manufacturers’ instructions.

### Synaptic vesicle trafficking assay

Synaptic vesicle trafficking assays were adapted from previous reports(*24*, *32*, *106*). For assessing synaptic vesicle uptake, iNSpheroids with and without iMGL were plated in PLOL-coated μ-Dish 35 mm with a glass bottom (ibidi, cat. no. 81158) and allowed to adhere for 2-3 hours in a humidified chamber at 3% O_2_, 5% CO_2_ and 37° C. The assay was divided into stages: (i) endocytosis and (ii) exocytosis. All solutions used during endocytosis assessment were ice cold and kept on ice unless otherwise stated. Cell culture supernatant was removed and cells were rinsed twice with DPBS, without Ca^2+^ and Mg^2+^. After rinsing, 10 μM FM 1-43FX (Invitrogen™, cat. no. F35355) diluted in HBSS, no calcium, no magnesium (Gibco™, cat. no. 14170120) was added and kept for 1 min on ice. Afterwards, cells were washed twice in HBSS and fixed with 4% paraformaldehyde and 4% sucrose at 4°C for 20 min. Cells were rinsed twice with ice-cold HBSS and stored in HBSS at 4 °C until further processing. Samples were visualized using 3D acquisition with a 63X objective and 0.5 μm of Z-stack thickness, on a Mica confocal microscope (Leica). Fluorescence images were processed and visualized using FIRE in the ImageJ software version 1.54j (https://imagej.net/ij/). p(k) was calculated to estimate the probability of occurrence of intensity value k (equation 1), being k a value within the intensity range.

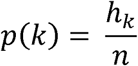

Where *h_k_*represents the intensity level and *n* the number of pixels in the ROI.

For exocytosis analysis, iNSpheroids with and without iMGL, were incubated with high potassium depolarizing solution (100 mM KCl buffer: 5 mM HEPES-NaOH, pH 7.4; 10 mM glucose; 2.5 mM calcium chloride; 1 mM magnesium chloride; 100 mM potassium chloride; 37 mM sodium chloride) for 5 min at 37 °C. Afterwards, iNSpheroids were incubated with 10 μM FM 1-43 dye (Invitrogen) in low potassium solution (5 mM KCl buffer: 5 mM HEPES-NaOH, pH 7.4; 10 mM glucose; 2.5 mM calcium chloride; 1 mM magnesium chloride; 5 mM potassium chloride; 37 mM sodium chloride) for 15 min at 37 °C. iNSpheroids were washed three times for 1 min with 5 mM KCl buffer with ADVASEP-7 (Sigma). Exocytosis was stimulated with 100 mM KCl buffer and samples were visualized live using a fluorescence microscope (Leica DMI6000) to monitor the fluorescence intensity decay over time. Fluorescence intensity was measured using the ImageJ software version 1.54j (https://imagej.net/ij/).

### Bulk RNA-sequencing

iMGL-iNSpheroids from both cell lines were harvested after three days of co-culture and sedimented by centrifugation at 300 xg, for 5 min. Cells were cryopreserved in a cell freezing medium, Cryostor CS10 (100-1061, STEMCELLTechnologies) and stored at −80°C. The frozen samples were shipped on dry ice to Genewiz (Leipzig, Germany) for RNA sequencing analysis. The RNA sequencing workflow after RNA isolation included initial Poly-A selection-based mRNA enrichment, mRNA fragmentation, and random priming with subsequent first- and second-strand complementary cDNA synthesis. End-repair 5’ phosphorylation and adenine nucleotide (dA)–tailing was performed. Lastly, adaptor ligation, polymerase chain reaction (PCR) enrichment, and Illumina NovaSeq technology– based sequencing with 2× 150–base pair (bp) read length were carried out. Sequence reads were trimmed to remove possible adapter sequences and nucleotides with poor quality using Trimmomatic v.0.36. The trimmed reads were mapped to the Homo sapiens GRCh38 reference genome available on ENSEMBL using the STAR aligner v.2.5.2b.

Unique gene hit counts were calculated by using *featureCounts* from the Subread package v.1.5.2. The hit counts were summarized and reported using the *geneID* feature in the annotation file. Only unique reads that fell within exon regions were counted. After the extraction of gene hit counts, the gene hit counts table was used for downstream analysis. TPM values for microglia gene targets were used and represent the number of a given transcript over one million full-length transcripts.

### Single-nuclei RNA-sequencing (sn-RNAseq)

Untransduced and rAAV9-mCherry transduced iMGL-iNSpheroids (derived from the iPSC(IMR90)-4 and iPSC(DF6-9-9T.B)cell lines) were analysed by single nuclei RNA-sequencing (sn-RNAseq). Samples were collected at 0-, 6- and 24-hours post-transduction, and sedimented by centrifugation at 300 xg, for 5 min. Cells were then resuspended and cryopreserved in cell freezing medium, CryoStor CS10 (STEMCELL Technologies, cat. no. 100-1061) and stored at −80 °C until shipment. Samples were sent for nuclei isolation and sequencing at Genewiz laboratories (Leipzig, Germany).

### Data processing and graph-based clustering

Raw data from snRNAseq was analysed and processed into a transcript count matrix using Cell Ranger (https://www.10xgenomics.com/, v6.1.1) from the Chromium Single Cell Software Suite by 10x Genomics. Fastq files were generated using the Cell Ranger *mkfastq* command with default parameters. Gene counts for each cell were quantified with the Cell Ranger count command with default parameters. For all analyses, the human (GRCh38) plus the mCherry genes were used as reference. The resultant gene expression matrix was imported into the R statistical environment for further analysis (RStudio 2024.04.2+764 “Chocolate Cosmos” Release). Ambient RNA removal was carried out using the *DecontX* methodology [104]. Cell filtering, data normalization, and clustering were carried out using the R package Seurat v5.0.1. Filtered feature-barcode matrices were used as input for the initial Seurat object, including only cells with at least 200 genes expressed in at least three cells. Datasets were further filtered to remove low-quality cells based on the following exclusion criteria parameters: (i) ratio of mitochondrial versus endogenous gene expression < 20% (putative dying cells); (ii) < 200 or > 5,000 total genes (putative poorly informative cells and multiplets). Data normalization, scaling, and identification of the top 3,000 most variable genes were performed using *SCTransform* v2.0 method, using default parameters and regressing out the percentage of genes coding for ribosomal proteins. This normalization method is claimed to outperform the classical log-normalization workflow, allowing the use of a higher principal component (PC) to capture more detailed biological variation for clustering. Principal Component Analysis (PCA) was run on each dataset and the first 30 PCs were selected for downstream analysis, based on their contribution to each PC variability (Elbow Plot). The overall significance of each PC and the corresponding p-value were calculated according to the *JackStraw* method. The different sample datasets were integrated into a single object using Seurat’s integration standard workflow, using the first 30 PCs, 3,000 variable features, and the *SCTransform* normalized data. Integration anchors were identified using reciprocal PCA and reference-based integration, with the timepoint 0 hours being used as reference. Dimensionality reduction was then performed with PCA on the batch-corrected data.

Uniform Manifold Approximation and Projection (UMAP) dimensionality reduction was performed on the calculated PCs to obtain a 2D representation for data visualization. To find the optimal clustering resolution, the *Clustree* package was used to compute different resolution values (0.1-1.0); the 0.1 resolution was selected for downstream analysis. Cell clusters were identified using the Louvain algorithm at resolution r = 0.1, implemented by the *FindCluster* function of Seurat. To characterize each cluster, a comprehensive manual annotation was performed. A list of marker genes from different cell types was collected from a literature-curated set of relevant marker genes (Table S2)(*107*, *108*).

### Sample preparation for mass spectrometry

iNSpheroids were harvested from the STBs and sedimented by centrifugation at 300 xg, for 5 min. The supernatant was discarded, and the resulting cell pellet was washed twice with PBS. Cells were lysed in Triton X-100 lysis buffer (50 mM Tris, 5 mM EDTA, 150 mM NaCl, 1% Triton X-100; all from Sigma-Aldrich), and 1x complete protease inhibitors cocktail (Roche), for 45 min at 4 °C, with periodic agitation. Total protein was quantified with the Micro BCA Protein Assay Kit (Thermo Fisher Scientific) following the manufacturer’s instructions. Proteins were extracted and precipitated in methanol-chloroform, as previously described (*21*). Briefly, one volume of protein solution and 4 volumes of methanol were centrifuged at 9 000 xg for 10 s, mixed with 2 volumes of chloroform, and centrifuged again. For phase separation, 3 volumes of water were added to the samples, which were homogenized by vigorous vortex, and centrifuged at 9 000 xg for 1 min. The upper phase was discarded, and 3 volumes of methanol were added.

Samples were gently mixed and centrifuged at 9 000 xg for 2 min to pellet precipitated protein. The supernatant was discarded, and the protein precipitate dried at 60 °C before solubilization in 0.15% of RapiGest SF Surfactant (Waters), overnight, at 4 °C, with continuous agitation; when protein precipitates were still detected, an additional step of solubilization was performed. Protein was quantified using the Bradford assay 36. For in-solution digestion, samples were reduced in 5 mM of dithiothreitol (DTT), for 30 min, at 60 °C; alkylated in 15 mM iodoacetamide (IAA), for 30 min in dark, and incubated at 100°C, 5 min; digested with trypsin (Promega; 1.2 μg/100 μg protein), at 37 °C, overnight. Trypsin was inactivated by acidification in 0.5% trifluoroacetic acid, at 37 °C, for 45 min. Samples were centrifuged at 16 000 xg for 10 min, and supernatants were collected into new tubes and dried using a Savant™ Universal Spe e dV ac concentrator (Thermo Fisher Scientific).

### Spectral library generation by information-dependent acquisition (IDA)

A total of 15 samples corresponding to timepoint 0-, 48- and 72-hours post-transduction, control and rAAV9-mCherry transduced, were subjected to information-dependent acquisition (IDA) analysis by Nano-liquid chromatography tandem mass spectrometry (nanoLC-MS/MS) analysis on an e kspert NanoLC 425 cHiPLC system coupled with a TripleTOF 6600 with a NanoSpray III source (Sciex, Framingham, MA, USA). Samples from the same condition and timepoint were pooled for subsequent analysis (N = 4).

Peptides were sprayed into the MS through an uncoated fused-silica PicoTip emitter (360 μm O.D., 20 μm I.D., 10 ± 1.0 μm tip I.D., New Objective, Oullins, France). The source parameters were set as follows: 15 GS1, 0 GS2, 30 CUR, 2.5 keV ISVF, and 100 °C IHT. A reversed-phase nanoLC-MS/MS with a trap and elute configuration, using a Nano cHiPLC Trap column (Eksigent, USA, 200 μm × 0.5 mm, ChromXP C18-CL, 3 μm, 120°Å) and NanoLC column (Eksigent, USA, 75 μm × 15 cm, ChromXP 3C18-CL-120, 3 μm, 120°Å) was performed. Water with 0.1% (v/v) formic acid (solvent A) and 0.1% formic acid in acetonitrile (solvent B) was used. The trap column was loaded with each sample at a flow rate of 2 μL/min for 10 min using 100% (v/v) solvent A Peptide separation was performed at 300 nL/min applying a gradient (v/v) of solvent B as follows: 0–1 min, 5%; 1–91 min, 5–30%; 91–93 min, 30–80%; 93–108 min, 80%; 108–110 min, 80–5%; and 110–127 min, 5%. Each sample pool was subjected to IDA using three different MS m/z ranges, which were calculated using the SWATH Variable Window Calculator V1.0 (Sciex, Framingham, US) based on a reference sample. was subjected to two IDA runs. The mass range for the MS scan was set to m/z 400–622.9, 621.9–790.9, and 789.9–2 000. The 50 most intense precursors were selected for subsequent MS/MS scans of 150–1,800 m/z, in high sensitivity mode, for 40 ms, using a total cycle time of 2.3

s. The selection criteria for parent ions included a charge state between +2 and +5 and counts above a minimum threshold of 125 counts per second Ions were excluded from further MS/MS analysis for 12 s. Fragmentation was performed using rolling collision energy with a collision energy spread of five. The spectral library was created by combining all IDA raw files using ProteinPilot software (v5.0 ABSciex) with the Paragon algorithm and with the following search parameters: Homo sapiens from Uniprot/SwissProt database; trypsin digestion; iodoacetamide cysteine alkylation; TripleTOF 6600 equipment; and biological modifications as ID focus. After a false discovery rate (FDR) analysis, only proteins with FDR < 1% were included in the reference spectral library.

### Protein quantification by sequential window acquisition of all theoretical fragment ion spectra-mass spectrometry (SWATH-MS)

Each sample was analysed in triplicate (3 μg per analysis) by sequential window acquisition of all theoretical fragment ion spectra (SWATH)-MS, using the instrument setup described for the IDA runs. The mass spectrometer was operated in a cyclic data-independent acquisition (DIA) as previously established (*21*). SWATH-MS data were acquired with the SWATH acquisition method, using a set of 64 overlapping variable SWATH windows covering the precursor mass range of 400–2 000 m/z. The SWATH variable windows were calculated using the SWATH Variable Window Calculator V1.0 (Sciex, Framingham, MA, USA) based on a reference sample. A 10 ms survey scan (400– 2 000 m/z) was acquired at the beginning of each cycle, and the subsequent SWATH windows were collected from 400 to 2 000 m/z for 50 ms, resulting in a cycle time of 3.26 s. Rolling collision energy with a collision energy spread of 5 was used. The spectral alignment and targeted data extraction of DIA samples were performed using PeakView v.2.2 (Sciex, Framingham, MA, USA), with the spectral library as reference. For data extraction the following parameters were used: Six peptides/protein, six transitions/peptide, peptide confidence level of >95% (corresponding to FDR < 1%), FDR threshold of 1%, excluding shared peptides, and extracted ion chromatogram (XIC) window of 6 min and width set at 20 ppm. Data were directly exported to Markerview 1.3.1 (Sciex, Framingham, MA, USA) and normalized using total area sums to obtain the final quantification values. A total of 3 239 proteins were quantified under these conditions.

### Proteome data analysis

Protein differential expression analysis was performed using the R package “DEP” (version 1.14.0)(*109*). Pairwise comparisons for TIC-stimulated versus control samples were performed for the 72-hour timepoint at thresholds of FDR < 0.1 and fold-change >1.5.

### Detection of AAV9-mCherry genomes by SABER-FISH technology in 2D iMGL

Twenty non-overlapping oligonucleotides that recognize only the negative-sense strand of AAV9-mCherry genome were retrieved using the ApE software as described by Wang and colleagues(*49*). All probes were analysed for specificity using the NCBI-blast software and quality using the *OligoAnalyzer* tool from integrated DNA technologies (IDT). The respective SABER-FISH probe sequences were obtained by appending to the 3’end a TTT linker followed by the initiator 9-nt sequence CATCATCAT for probe extensions using a catalytic hairpin. Probes, catalytic hairpin, the fluorescent oligonucleotide for detection of hybridized probes to AAV genome by in-situ hybridization (ISH), and the clean. G sequence (CCCCGAAAGTGGCCTCGGGCCTTTTGGCCCGAGGCCACTTTCG), used in the probe extension reactions, were ordered from IDT in IDTE (10 mM Tris, 0.1 mM EDTA) pH 7.5. Probes and clean.G sequences were order at 25 nM scale with standard desalting. The catalytic hairpin and fluorescent oligonucleotide (100 µM and 250 µM concentration, respectively) were HPLC purified. Sequences of all probes, catalytic hairpin and fluorescent oligonucleotide can be found in Table S4.

Probes were extended individually to 550-750 nucleotide, comprising long arrays of binding sites for the fluorescent oligonucleotide, through the primer-exchange reaction cycle (at 37 °C for 4h) essentially as described by Kishi et al. (*48*)and on the online available protocol at http://saber.fish/. The size of extended probes was monitored by gel agarose. Extended probes were pooled and subsequently purified and concentrated using a MinElute kit (Qiagen, cat. no. 28004) and eluted in nuclease-free water.

Detection of AAV9-mCherry genomes was also performed as described previously (*48*), with minor modifications. All chemicals were purchased from Sigma-Aldrich unless otherwise indicated. Untransduced and rAAV9-mCherry transduced iMGL recovered the supernatant of iMGL-iNSpheroid co-culture (24-hours post-transduction) were loaded in 8-well Ibidi chamber slides (Ibidi, cat. no. 28004), rinsed three times with 1xPBS (250 μL) to remove extracellular AAVs, and immediately fixed in 4% (vol/vol) PFA plus 4% (vol/vol) sucrose (200 μL) for 20 min at RT. Afterwards, cells were washed in 1x PBS (2 x 1 min), permeabilized in 1x PBS with 0.5% (vol/vol) Triton X-100 (200 μL; cat. no.

<u>9036-19-5</u>) for 15 min at RT. After washing in 1x PBS with 0.1% (vol/vol) Tween-20 (cat. no. 8.22184.0500) (1x PBST) (250 μL, 1x 1 min and 1 x 2 min at RT), samples were incubated in 0.1% HCl (5 min, RT) and washed twice in 2x SSC (cat. no. S6639) solution with 0.1% (vol/vol) Tween-20 (2x SSCT) (1x 1 min and 1x 2 min at RT). Afterwards, samples were washed with fresh 50% formamide (cat. no. S4521) + 2× SSCT solution (200 μL) for 5 min at RT followed by incubation in the same fresh solution at 60 °C for 2 h. Samples were then incubated in freshly prepared hybridization solution (50% formamide + 10% dextran sulfate (cat. no. D8906) + 1% tween-20 + 2x SCC) without and with 100 mM purified pooled probes for 22 h at 42 °C (200 µL), followed by four 5-min washes with 2× SSCT solution at 60 °C and two 2-min washes in the same solution at RT. After washing once in 1× PBS at RT, samples were incubated with 0.5 µM fluorescent oligonucleotide (200 µL) at 37 °C for 1h30 min, and subsequently washed three times with 1x PBS at 37 °C (once for 5 min and twice for 2 min) and once with PBS at RT. Next, nuclei were labelled by staining with DAPI as described above. Images were acquired on a Zeiss LSM880-point scanning confocal microscope controlled with the Zeiss Zen 2.3 (black edition) software. Images were processed using ImageJ software and only linear manipulations were performed.

### Statistical analysis

Data are expressed as mean ± standard deviation (S.D.), with statistical analyses conducted in GraphPad Prism v9.0.1 (GraphPad Software, San Diego, CA). When comparing only two experimental groups, the unpaired Student t-test was used for data with normal distribution; if otherwise, the Mann-Whitney test was used. When comparing three or more groups, a one-way analysis of variance (ANOVA) followed by the Bonferroni or Tukey post hoc test was used for data with normal distribution. To compare different groups with two independent variables, we used a two-way ANOVA followed by the two-stage linear step-up procedure of Benjamini–Hochberg method for statistical significance was considered for p<0.05.

## Acknowledgments

The authors acknowledge Dr. João Relvas and Dr Ulrike Kleymann for the useful scientific discussion.

The authors acknowledge the IMI-ARDAT consortium (www.ardat.org) for the funding, services, and scientific input.

Illustrations were created in biorender.com.

## Author contributions

Conceptualization: CMG, CB Methodology: CMG, GS, CB

Investigation: CMG, GS, MMA, SJH, DDL, PH, RJ, RJN

Visualization: CMG, GS, MMA, DS, SH, DDL, PH, RJ, RJN, LPA, MT, PMA, CB

Supervision: CB Writing—original draft: CMG, CB

Writing—review & editing: CMG, GS, MMA, DS, SH, DDL, PH, RJ, LD, RJN, LPA, MT, PMA, CB

Funding acquisition: CB, PMA, RJN, PH, MT Resources: CB, PMA, RJN, PH, LD, MT

## Data and materials availability

- Single-nuclei and bulk RNA-seq data have been deposited at GEO: Bulk RNAseq data accession number GSE299019; and password: ktupwuqodncrpar. snRNAseq data accession number: GSE298806; and password: qzatigmupbsnfgt.
- The mass spectrometry proteomics data have been deposited to the ProteomeXchange Consortium via the PRIDE (*110*) partner repository with the dataset identifier PXD065639. For revision purposes the dataset can be accessed by logging in to the PRIDE website using the following account details:

**Username:**; **Password:** 2SNcwrJ0778b.

