## Supplemental Figures and Tables for "Innate immunocompetent iNSpheroids: A hiPSC-derived 3D model to study the central nervous system captures an early CNS response to rAAV"

Catarina Monteiro Gomes *et al.*

**This PDF file includes:**

Supplementary Text

Figs. S1 to S12

Tables S1 to S4

Supplementary Text

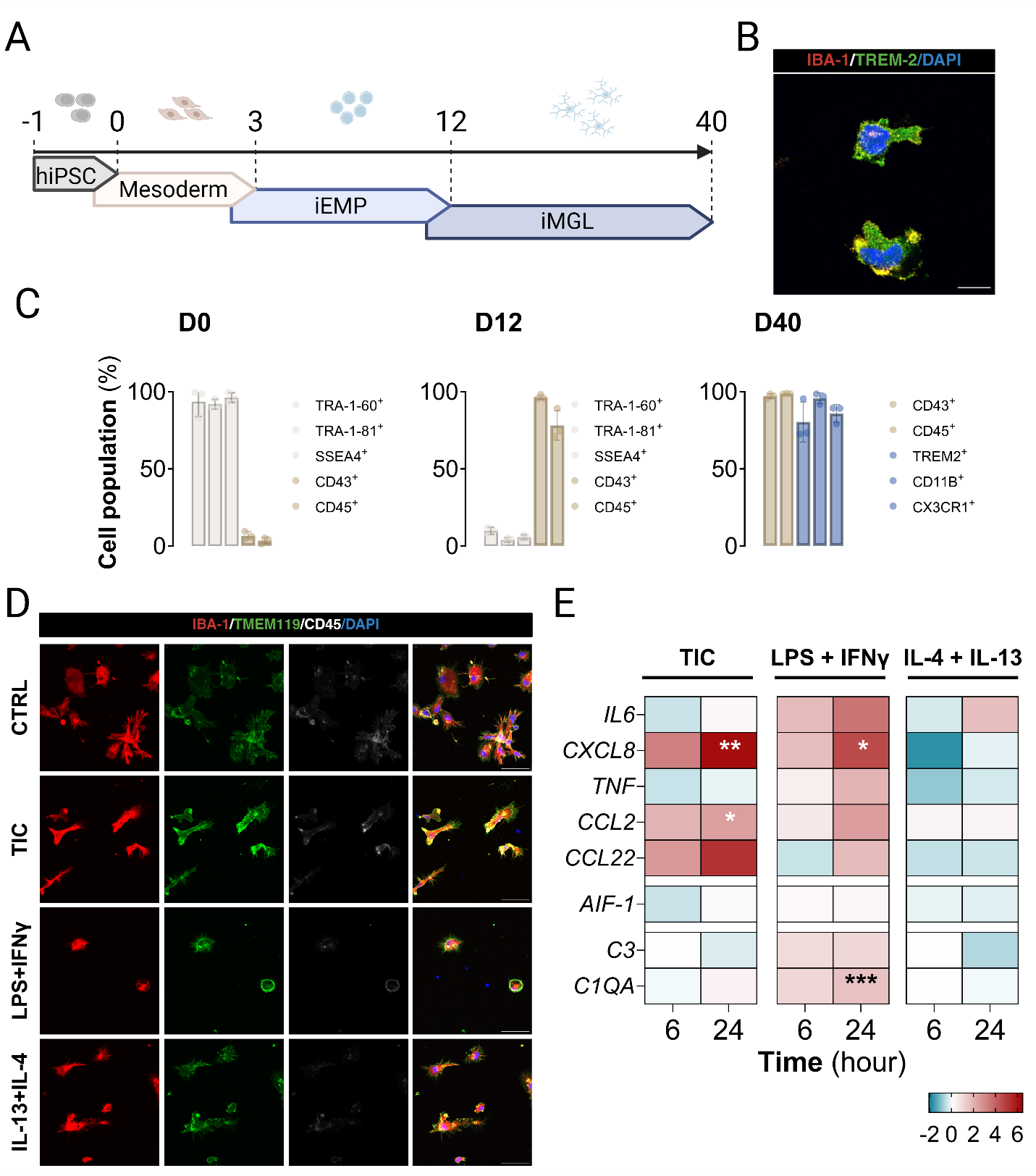

Fig. S1. Characterization of 2D hiPSC-derived microglia (iMGL) differentiation.

(a) Schematic overview of experimental setup. Briefly, human induced pluripotent stem cells (hiPSC) are harvested and replated as small colonies on day -1. Afterwards, a stepwise approach is used to induce mesoderm, iMGL progenitors, and, finally, iMGL by changing the medium feed formulation. After 40 days of differentiation, (b) iMGL express the IBA-1 and TREM-2 proteins. Scale bar = 5 𝜇m. (c) Flow cytometry characterization at D0 (pluripotency marker proteins), D12 (iMGL progenitor marker proteins) and D40 (microglia marker proteins). To ascertain iMGL functionality, cells were stimulated with three different inflammatory stimuli: (i) TNF-𝛼, IL-1𝛼 and C1q, collectively known as TIC; (ii) LPS plus IFN-𝛾; and (iii) IL-13 plus IL-4. iMGL were then characterized by their (d) morphology and IBA-1, TMEM119, and CD45 expression after 48 hours of stimuli, and (d) proinflammatory gene expression profile was analysed by RT-qPCR after 6 and 24 hours. Results were calculated using the comparative cycle threshold values method (2−ΔΔ𝐶𝑡). Gene expression values were normalized to the expression of housekeeping genes HPRT1 and RPL22 and calculated as fold change over non-stimulated control cells. Data are represented as the mean of four independent experiments. Statistical analysis was performed by applying the one-way ANOVA test, *p < 0.05, **p < 0.01, ***p < 0.001.

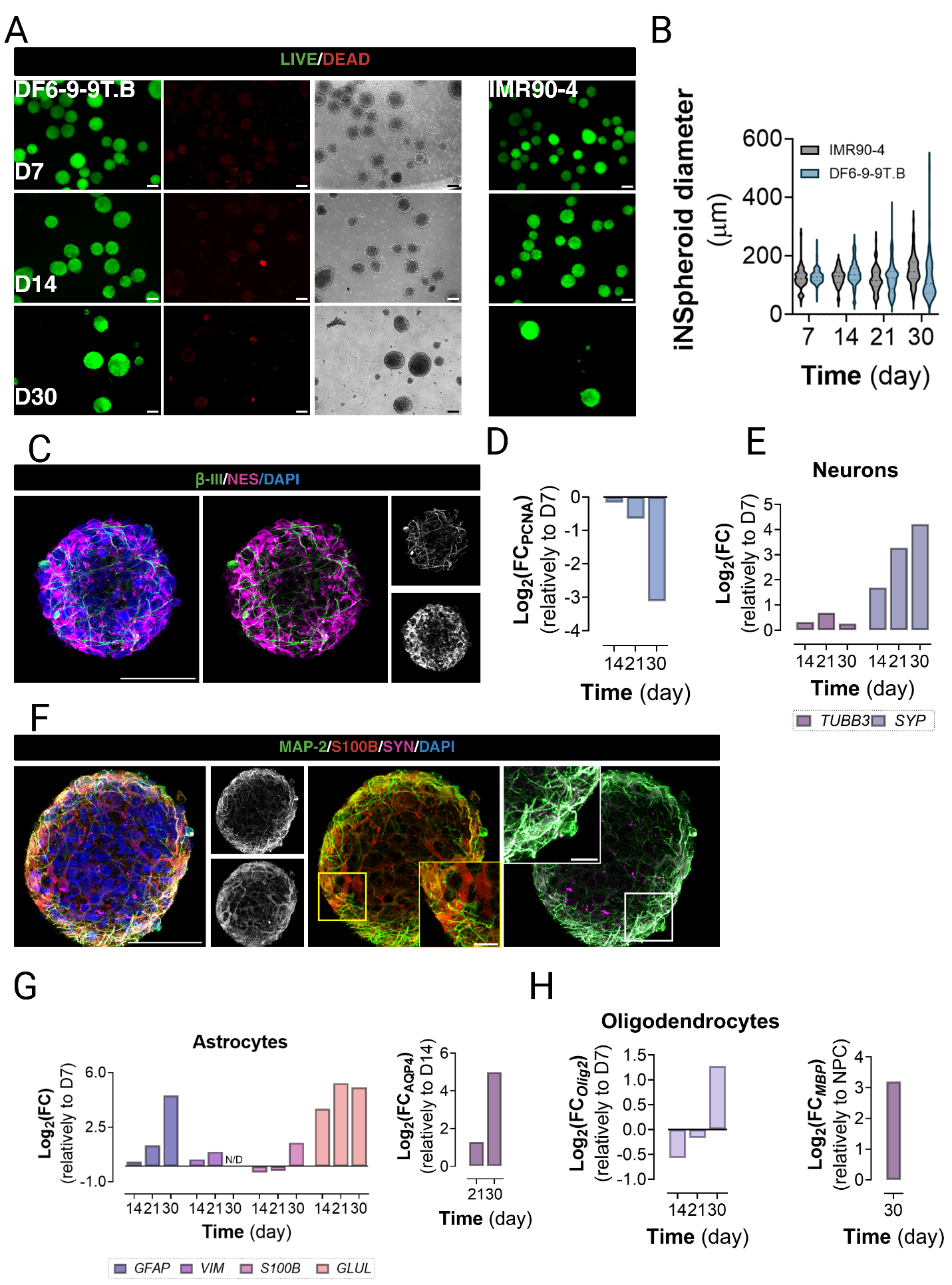

Fig. S2. Phenotypic characterization of iNSpheroid differentiation.

(a) Representative live/dead staining images of hiPSC-derived neurospheroids (iNSpheroids) generated from the iPSC(DF6-9-9T.B) (in short DF6-9-9T.B) line in comparison to the iPSC(IMR90)-4 (in short IMR90-4) line, at days 7, 14, and 30 of differentiation. Green indicates the metabolization of Fluorescein Diacetate (FDA) in live cells, while red indicates propidium iodide (PI) binding to DNA bases in dead cells. Scale bars represent 200 𝜇m. (b) Violin plot of iNSpheroid diameter measurements across differentiation the indicated time points for DF6-9-9T.B and IMR90-4, showing consistent growth and size distribution. (c) Immunofluorescence detection of bIII-tubulin (TUBB3) (green, early neuronal differentiation marker) and nestin (NES, magenta, neural stem cell marker), at day 7 of culture. Scale bar: 50 𝜇m. Representative images of iPSC(DF6-9-9T.B) line. (d) RT-qPCR analysis of proliferating cell nuclear antigen (PCNA) and (e) TUBB3 and synaptophysin (SYP). Gene expression values were normalized to the expression of housekeeping genes HPRT1 and GAPDH in the same samples. Data are presented as fold change relative to day 7. (f) Immunofluorescence microscopy detection of microtubule-associated protein 2 (MAP-2, green, marker of the somatodendritic compartment of mature neurons), S100 calcium-binding protein B (S100B, red, astrocytic maturation marker), and synaptophysin (SYN, magenta, presynaptic vesicle marker in neurons). Scale bar: 50 𝜇m; 10 𝜇m for zoom-in inset. RT-qPCR analysis of (g) glial fibrillary acidic protein (GFAP), S100B, glutamate-ammonia ligase (GLUL) and vimentin (VIM), and (h) basic helix-loop-helix (bHLH) transcription factor encoded by the OLIG2 gene and myelin basic protein (MBP) gene expression. Values were normalized to the expression of housekeeping genes HPRT1 and GAPDH. All qPCR results were derived from the hiPSC(DF9-9-9T.B) line (N=1).

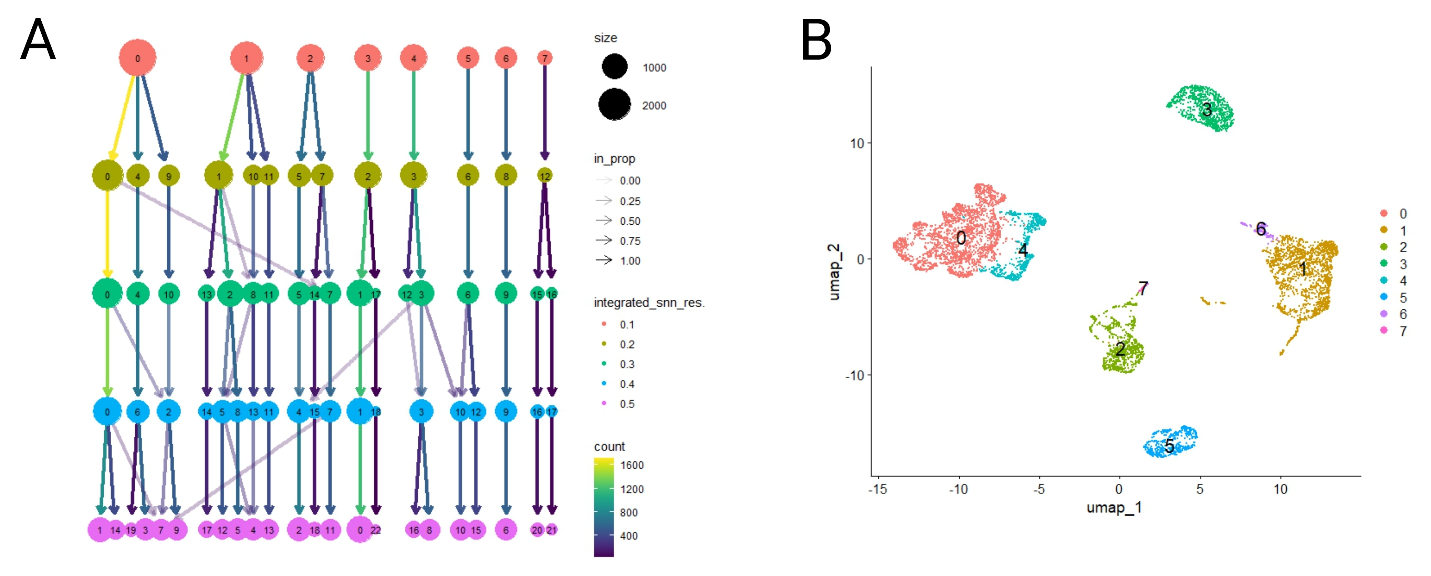

Fig. S3. Clustering and annotation of the main neural cell types and microglia in the integrated dataset of two hiPSC lines.

(a)Analysis of cluster stability across several resolution values with the Clustree package revealed that most clusters were stable at the 0.1 resolution. (b) Clustering the integrated datasets at resolution 0.1 allowed identifying seven major clusters.

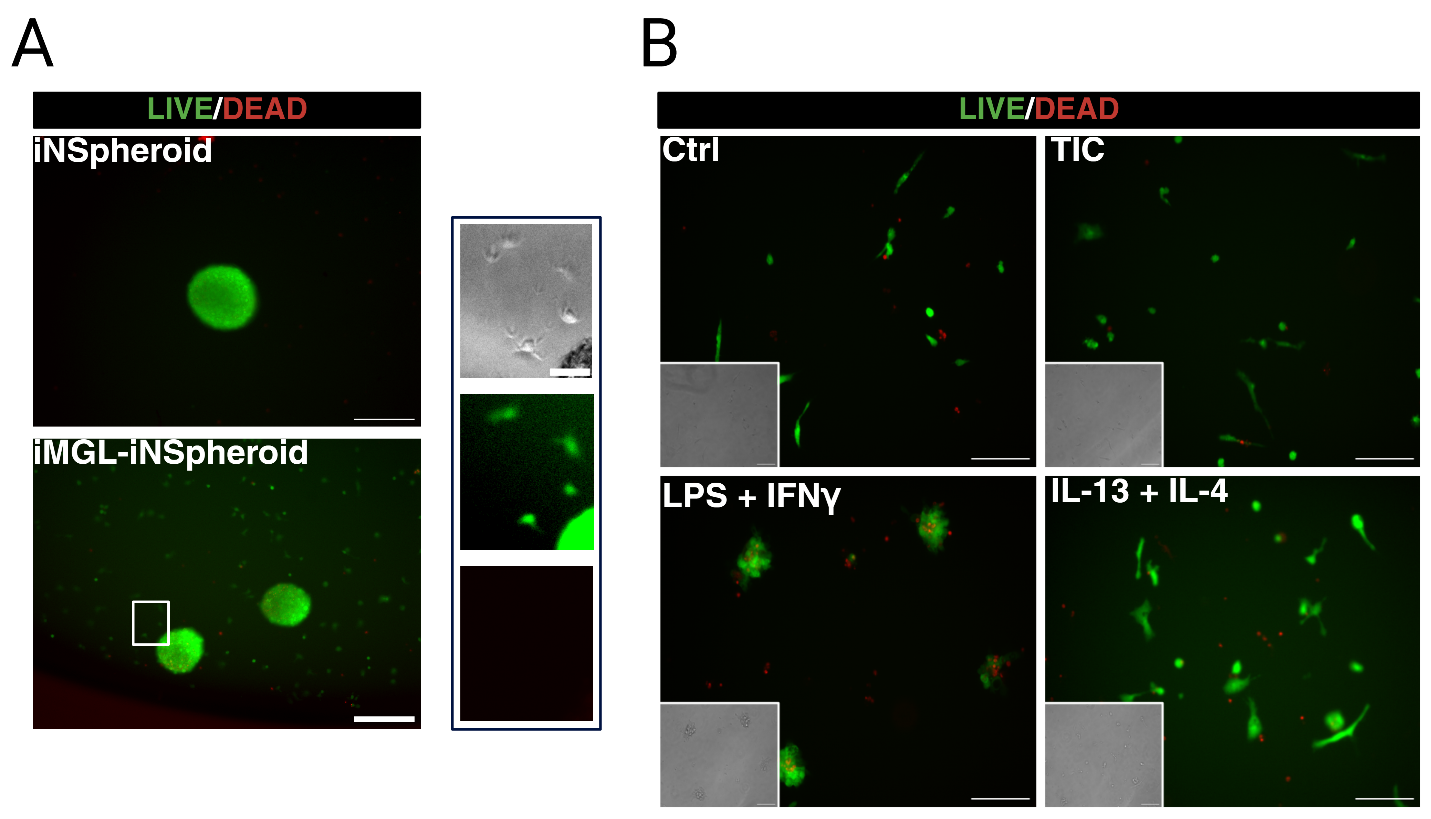

Fig. S4. iMGL and iNSpheroid maintain high cell viability after three days of co-culture and seven days of prototypical inflammatory challenge.

(a) Representative live/dead images of iPSC DF6-9-9T.B-derived iNSpheroids, without or with iMGL (iMGL-iNSpheroids), after three days of co-culture. Zoom-in image illustrating iMGL in the supernatant of the iMGL-iNSpheroid co-culture. Scale bar, 100 (left image) and 10 𝜇m (right panel, zoom-in image). (b) IMGL in suspension remain viable after seven days of treatment with pro- and anti-inflammatory stimuli. INSpheroids and iMGL were stained with fluorescein diacetate (FDA, live cells, green) and propidium iodide (PI, dead cells, red). Scale bar, 50 𝜇m.

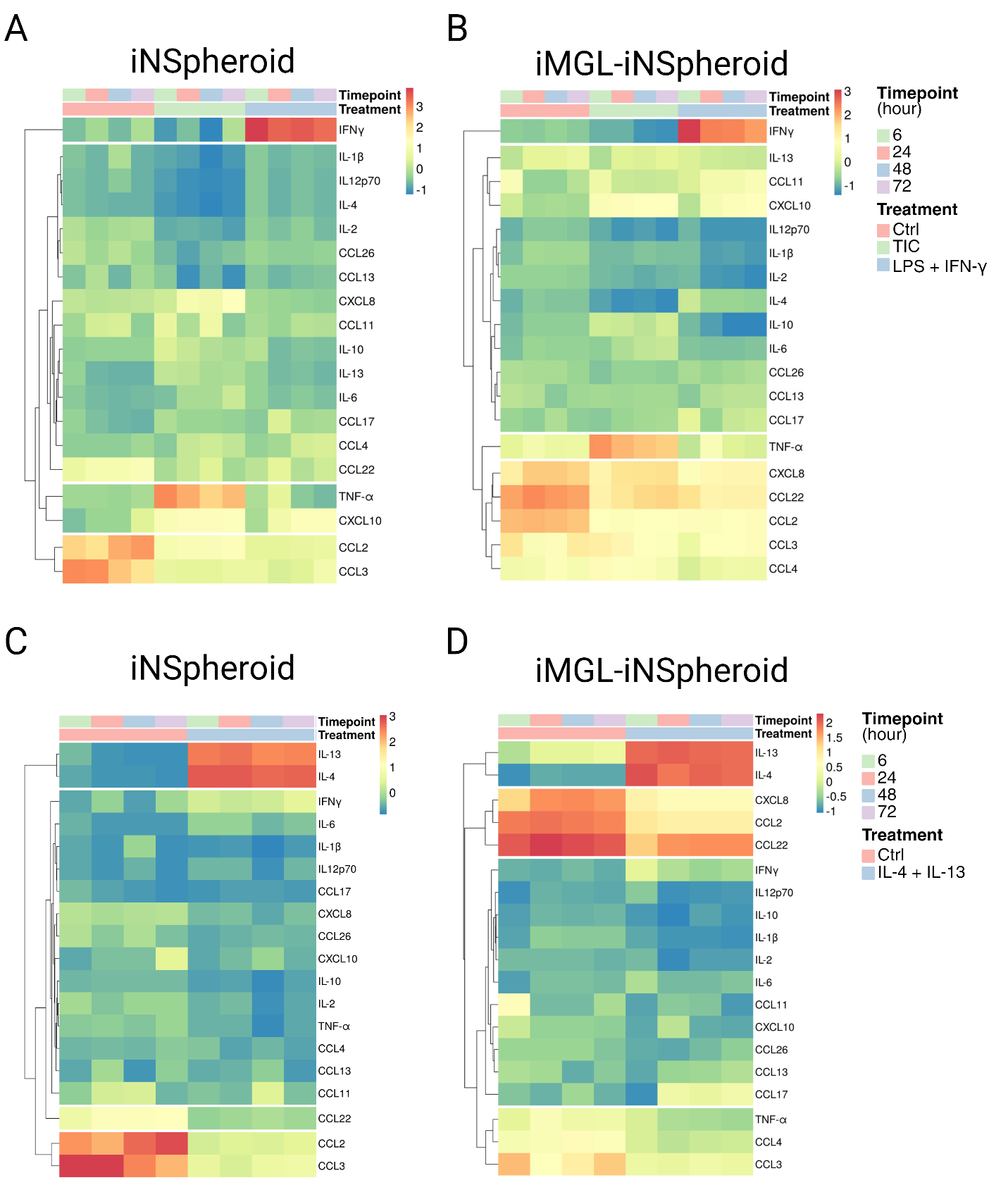

Fig. S5. iNSpheroids, with and without iMGL, mount a neuroinflammatory response to prototypical inflammatory stimuli.

Hierarchical clustering of the protein targets identified by secretome analysis of iNSpheroids (a) without and (b) with iMGL (iMGL-iNSpheroids) in the control condition and in response to TIC and LPS plus IFN-𝛾; and upon (c, d) anti-inflammatory stimuli (IL-4 plus IL-13), (c) without and (d) with iMGL. N=5, the data corresponds to two hiPSC lines (iPSC(IMR90)-clone 4 and IPSC(DF6-9-9T.B)).

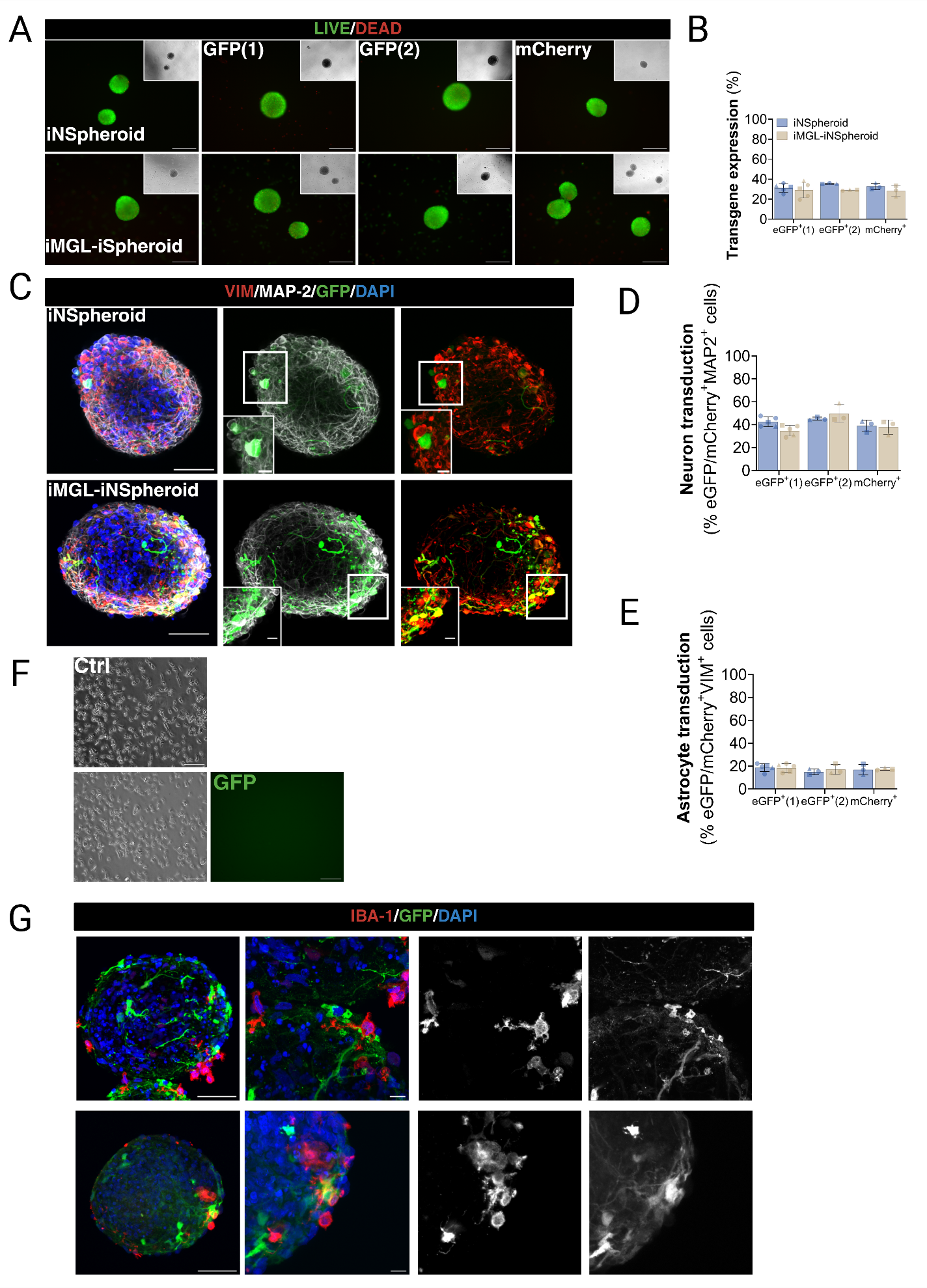

Fig. S6. rAAV9s transduce neural cells in the iMGL-iNSpheroids, but not iMGL.

(a) Representative live/dead images of iNSpheroids without and with iMGL (iMGL-iNSpheroids). INSpheroids were stained with fluorescein diacetate (FDA, live cells, green) and propidium iodide (PI, dead cells, red). Scale bar, 100 𝜇m. Quantification of (b) transgene-expressing cells in iNSpheroids and iMGL-iNSpheroids. Within the transgene-positive population, the tropism was analysed by assessing (c) immunofluorescence images of iNSpheroids and iMGL-iNSpheroids transduced with rAAV9 (GFP-expressing cells, green), MAP2-positive neurons (gray) and VIM-positive astrocytes (red). (d) Neuronal and (e) astrocytic tropism was quantified in the outer layer (40%) of the iNSpheroid structure, to account for technical artifacts. Each data point corresponds to one independent experiment. Results are expressed as mean ± S.D., in percentage. N > 3. iMGL transduced with rAAV9-GFP does not express the viral transgene, neither as a (f) 2D monoculture or (g) 3D iMGL-iNSpheroid co-culture. (f) Representative phase contrast and GFP images of iMGL from iPSC(IMR90) clone 4. Scale bar, (f) 100 𝜇m and (g) 50 𝜇m.

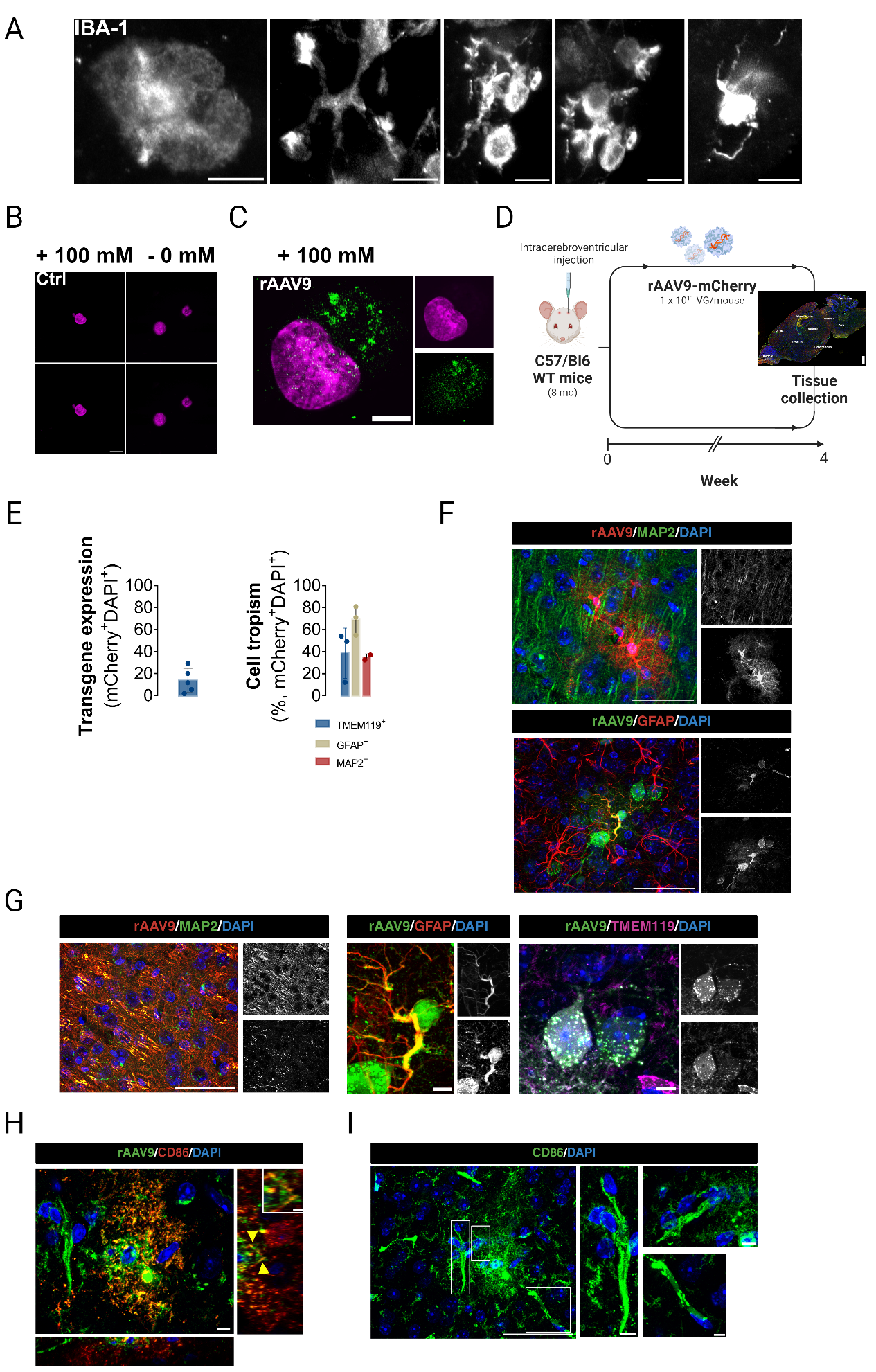

Fig. S7. iMGL are transduced but do not express rAAV9 transgene.

(a) Morphofunctional plasticity of IBA-1(grey)-positive iMGL, illustrated by representative images of the different morphologies observed in iMGL-iNSpheroids transduced with rAAV9-mCherry. Representative images of cultures differentiated from iPSC(IMR90)-clone 4. (b, c) Visualization of AAV9-mCherry genomes by SABER-FISH technology in untransduced and transduced 2D iMGL in-situ hybridized with no probes or 100 nM probes specific for AAV9-mCherry genome. The probes hybridization is represented in green, and nuclei are counterstained with DAPI, in magenta. Scale bar = 5 μm. (c) Highlight of iMGL transduced with rAAV9-mCherry after 24 hours. Scale bar = 2 μm. (d) Schematic illustration of C57/Bl6 wild-type mice model transduction with rAAV9-mCherry and tissue harvest, and processing. (e) Quantification of transduced cells in the tissue sections of mouse brain (left panel); and microglia (TMEM119-positive), astrocytic (GFAP-positive), and neuronal (MAP2-positive) (cell tropism, right panel). Quantification was performed on tissue sections of 5 different mice. Each data point corresponds to one mouse. Results are expressed as mean ± S.D., in percentage. N > 3. (f) Immunostaining of MAP-2 neuronal network (green, upper panel) and GFAP-positive astrocytes (red, bottom panel), in combination with mCherry labelling (red, top panel, or green, bottom panel). Maximum Z-projection of 0.21 μm optical slice thickness. Scale bar = 50 μm. (g) Highlight of neurons (left panel, green), astrocytes (middle panel, red), and microglia (right panel, magenta) transduced with rAAV9-mCherry. Maximum Z-projection of 0.21 μm optical slice thickness. Scalebar = 50 (left panel) and 5 μm (middle and right panels). (h) Immunofluorescence imaging representing close proximity between CD86-positive microglia (red) and rAAV9-transduced cells (green) in C57/Bl6 wild-type mice brain tissue slices. Juxta-position between iMGL and rAAV9-transduced cells can be observed in the orthogonal slices. Scalebar = 10 and 2 μm (zoom-in panel). (i) Morphofunctional plasticity of CD86(green)-positive microglia. Scale bars = 50 and 2 μm (zoom-in panel).

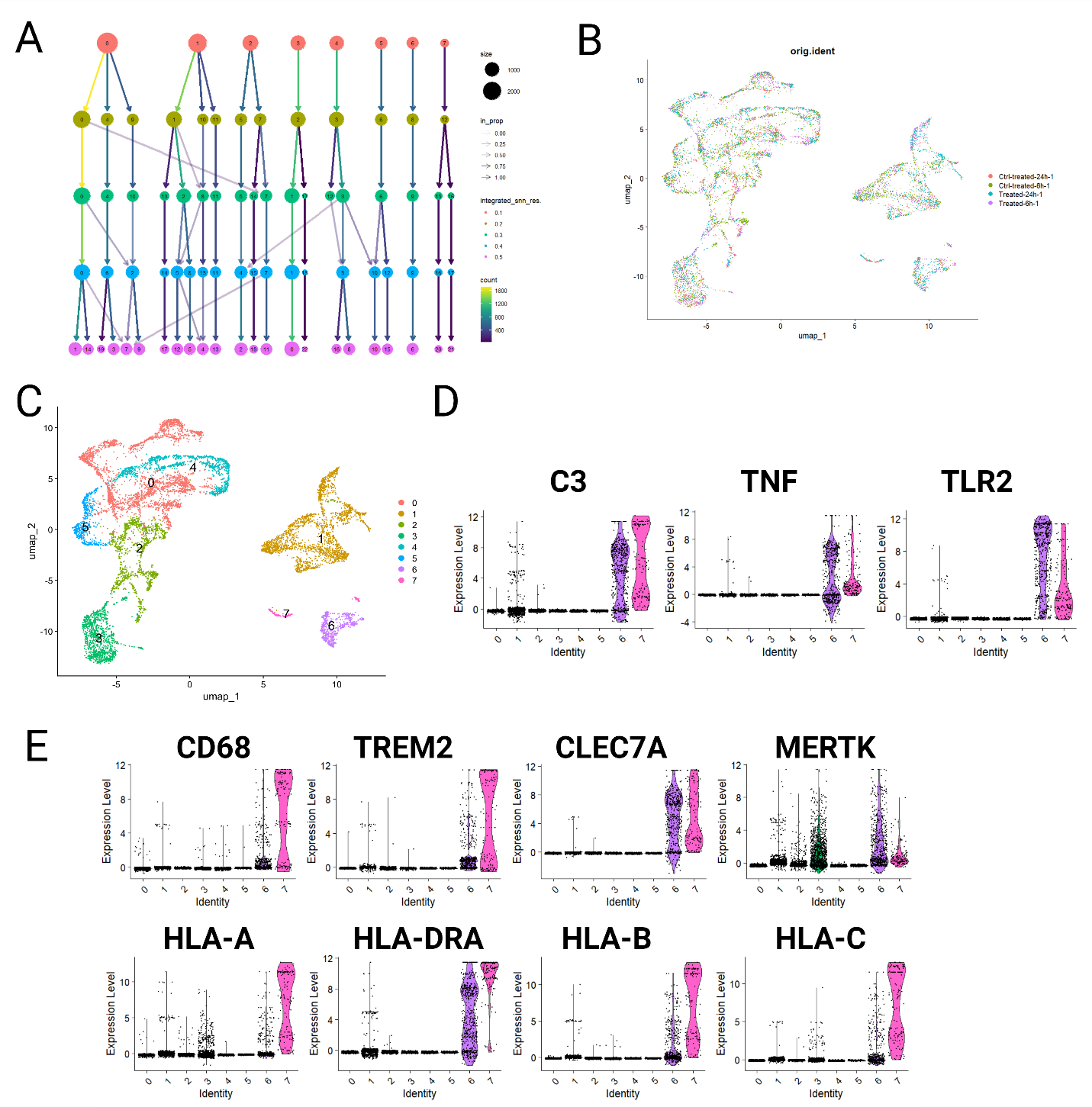

Fig. S8. Clustering and annotation of the main neural cell types and microglia in the integrated dataset of iNSpheroids transduced with rAAV9-mCherry.

(a) Analysis of cluster stability across several resolution values with the Clustree package revealed that most clusters were stable at the 0.1 resolution. (b, c) UMAP visualization of the integrated datasets (b) before and (c) after clustering at a resolution of 0.1, allowed for the identification of eight major clusters. Violin plot representation of (d) inflammation and (e) phagocytosis (upper panel), and major histocompatibility complex-associated genes (lower panel) amongst the different cell clusters identified in (c).

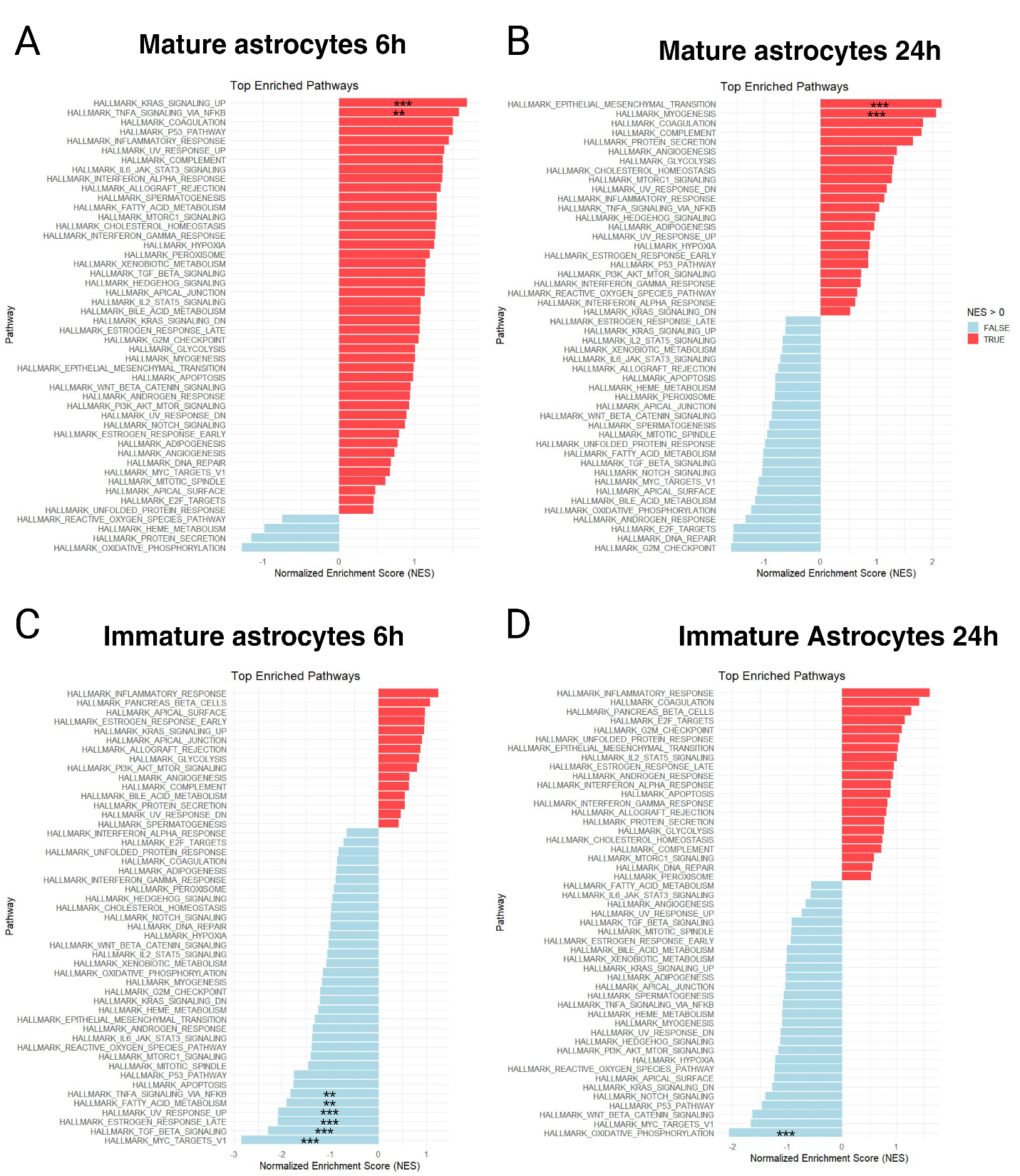

Fig. S9. snRNAseq of rAAV9-transduced iMGL-iNSpheroids reveals different inflammatory patterns between "mature" and "immature" astrocytic populations, at both 6- and 24-hours post-transduction.

Bar plot visualizing the enriched GSEA terms in iMGL-iNSpheroid cultures transduced with rAAV9-mCherry, over non-transduced controls, against the Hallmark gene set (Molecular Signatures Database, human MSigDB- GSEA). Samples were collected at two different timepoints: 6- and 24-hours post-transduction (hpt). Top enriched pathways in sub-cluster (a) mature astrocytes (cluster 6) 6hpt, (b) mature astrocytes 24hpt, (c) immature astrocytes 6hpt, and (d) immature astrocytes 24hpt are illustrated. GSEA was performed on Log2 fold change pre-ranked gene lists obtained from gene expression levels of the different cell types. NES, normalized enrichment score; *, adjusted p<0.05; **, adjusted p<0.01; ***, adjusted p<0.001.

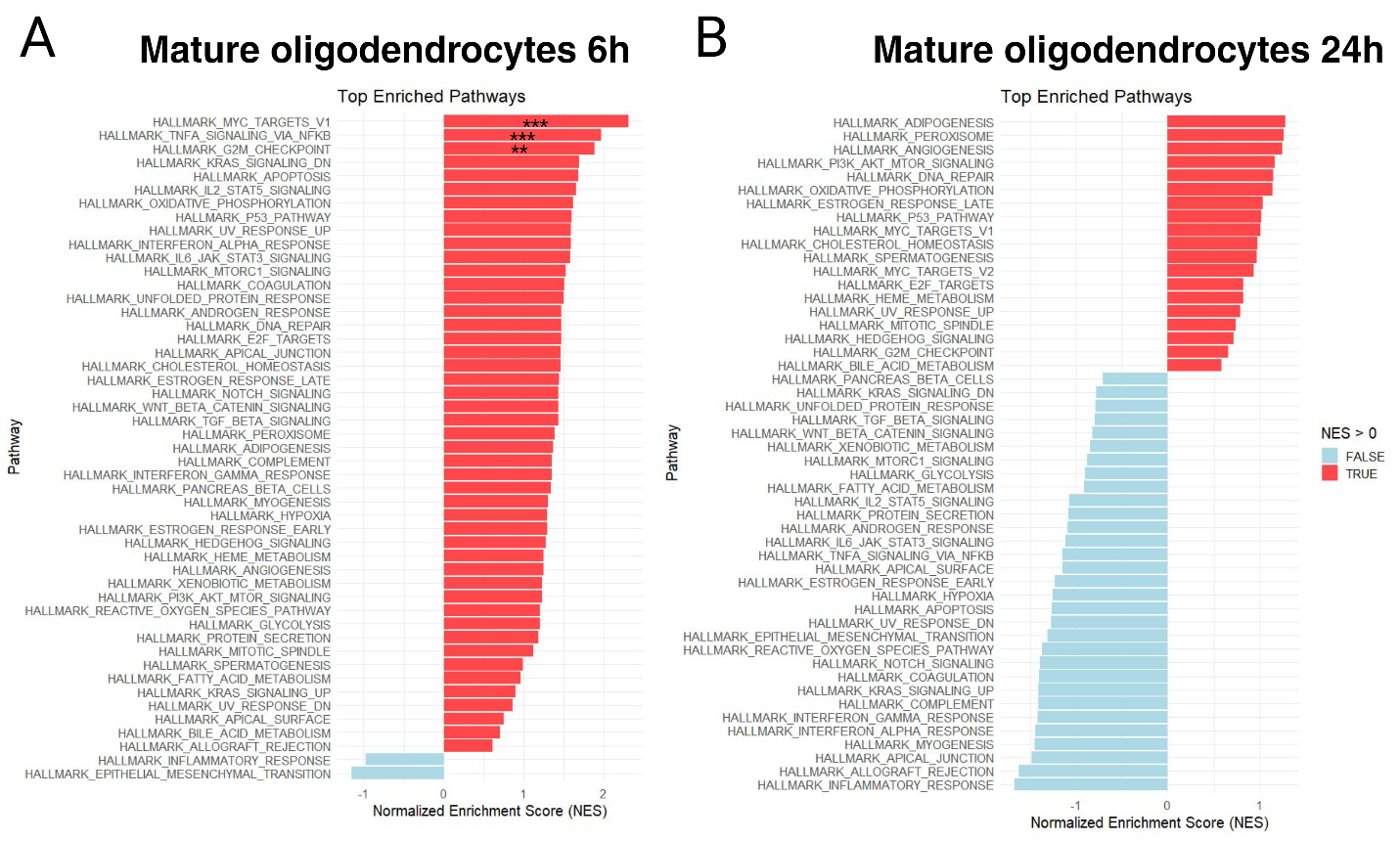

Fig. S10. snRNAseq of rAAV9 transduced co-cultures (iMGL-iNSpheroids) reveals different inflammatory patterns between "mature" and "immature" oligodendrocyte populations, at both 6- and 24-hours post-transduction.

Bar plot visualizing the enriched GSEA terms in iMGL-iNSpheroid cultures transduced with rAAV9-mCherry, over non-transduced controls, against the Hallmark gene set (Molecular Signatures Database, human MSigDB- GSEA). Samples were collected at two different timepoints: 6- and 24-hours post-transduction (hpt). Top enriched pathways in sub-cluster (a) oligodendrocytes at 6hpt and (b) 24hpt are illustrated. GSEA was performed on Log2 fold change pre-ranked gene lists obtained from gene expression levels of the different cell types. NES, normalized enrichment score; *, adjusted p<0.05; **, adjusted p<0.01; ***, adjusted p<0.001.

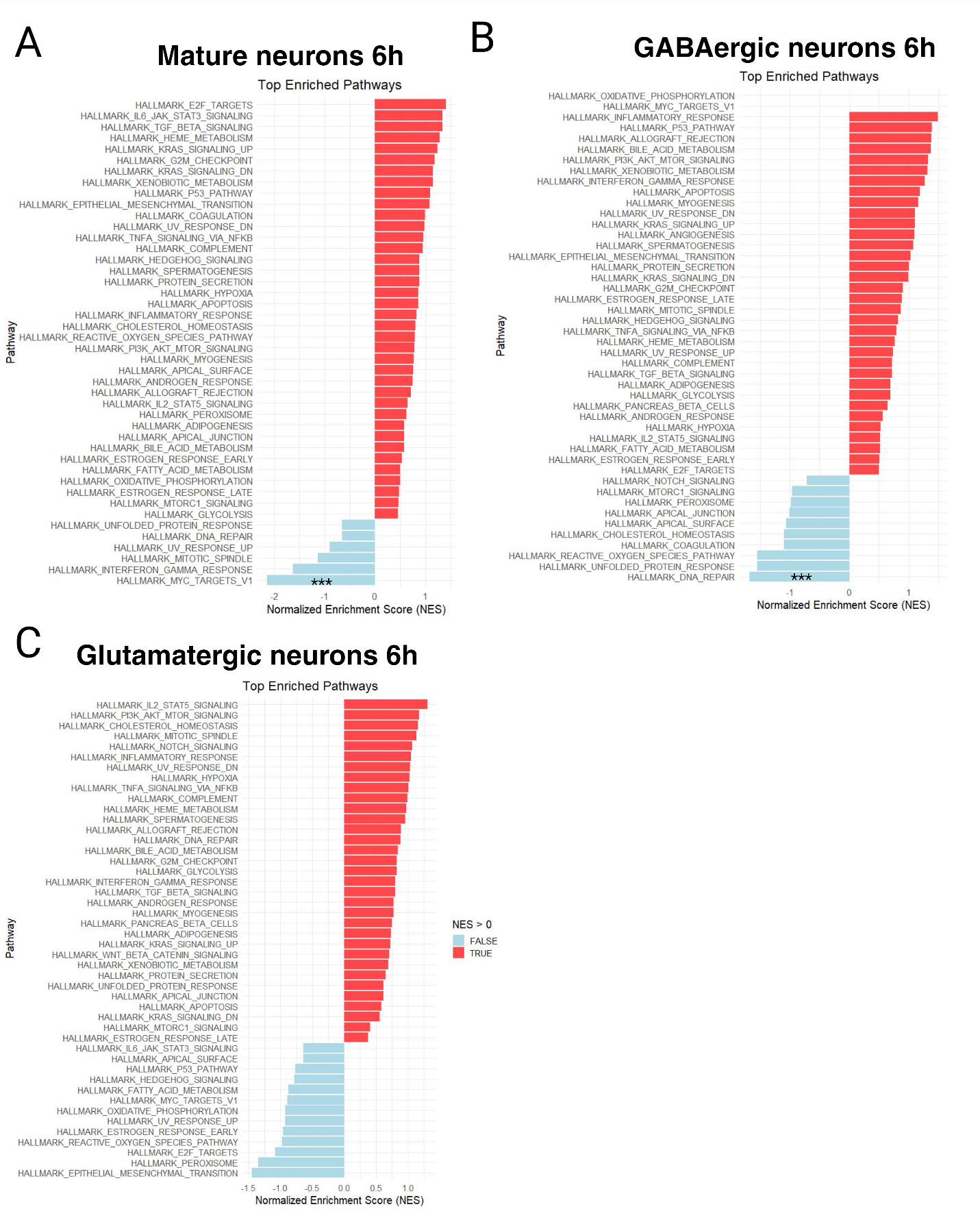

### Fig. S11. snRNAseq of rAAV9 transduced co-cultures (iMGL-iNSpheroids) reveals different inflammatory patterns in mature neurons, at 6-hours post-transduction.

Bar plot visualizing the enriched GSEA terms in iMGL-iNSpheroid cultures transduced with rAAV9-mCherry, over non transduced controls, against the Hallmark gene set (Molecular Signatures Database, human MSigDB- GSEA). Samples were collected at 6-hours post-transduction. Top enriched pathways in sub-cluster (a) mature neurons, (b) GABAergic neurons, and (c) glutamatergic neurons are illustrated. GSEA was performed on Log2 fold change pre-ranked gene lists obtained from gene expression levels of the different cell types. NES, normalized enrichment score; adjusted p<0.01; ***.

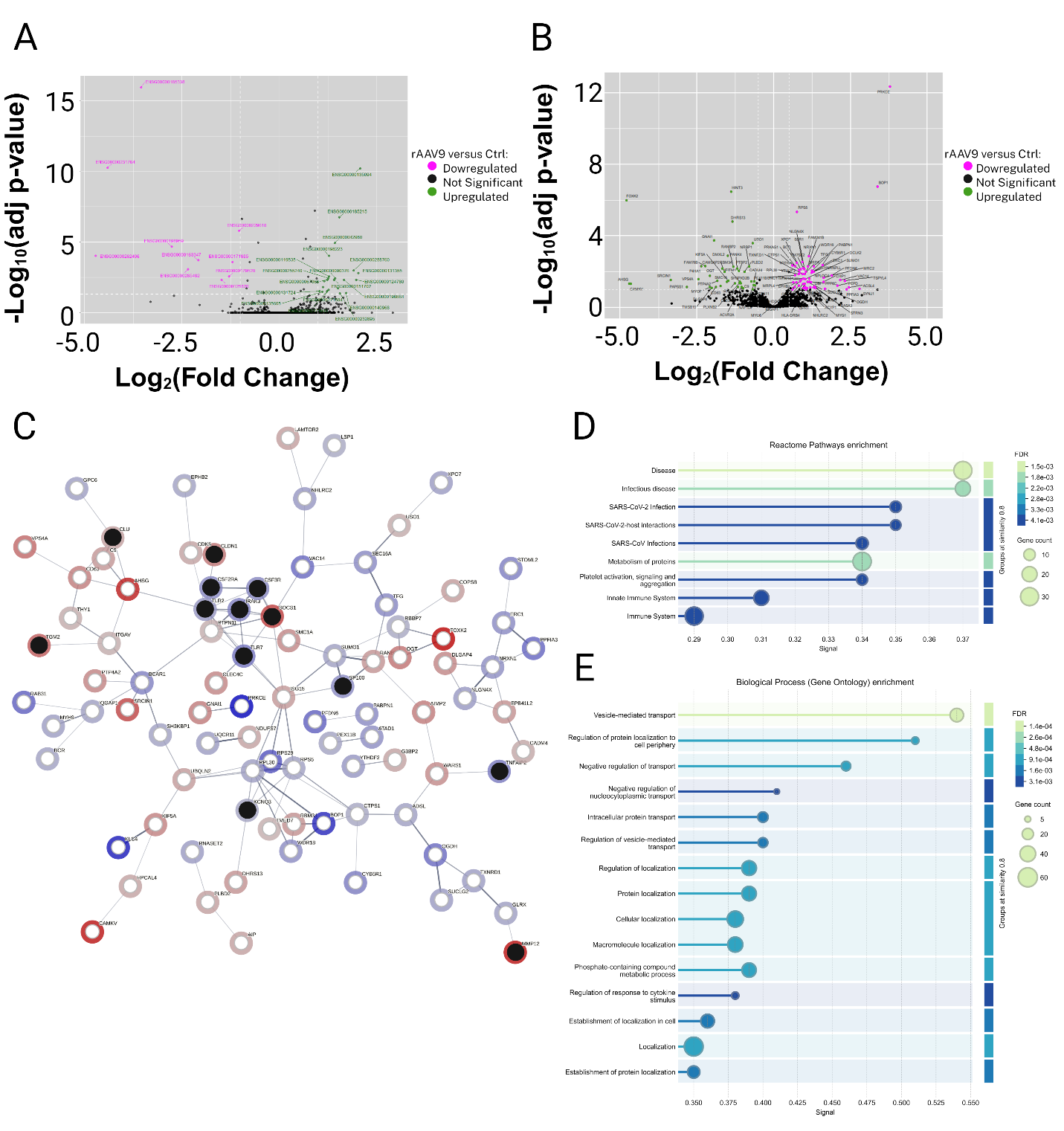

### Fig. S12. Correlation between pseudobulk snRNAseq and whole proteome analysis highlighting the expression of genes and proteins involved in the innate immune response to rAAV9 transduction.

Volcano plot of (a) pseudobulk analysis of snRNAseq data, merging the overall effects of the response at 6- and 24-hours post-transduction, and (b) proteins identified as differently expressed (DEP) in the iMGL-iNSpheroids untransduced and transduced with rAAV9-mCherry, 48 hours post-transduction. Significantly enriched genes and proteins on rAAV9-mCherry transduced iMGL-iNSpheroids are represented as magenta dots; while genes and proteins enriched in the control are green dots. (c) STRING analysis shows the correlation between genes (black bubbles) and proteins (hollow bubbles) that are involved and predicted protein-protein interactions. The network nodes represent the differentially expressed genes and proteins that have known interactions. Different connecting line thicknesses represent the degree of evidence used to predict associations. (d, e) Lollipop graph representing pathway enrichment analysis obtained using (d) the reactome pathway database enrichment and (e) the biological process gene ontology (BP-GO) database. Colour code represents the FDR value, while the bubble size refers to the number of proteins within the identified pathway.

**Table S1.**

Primer sequences used for reverse transcriptase quantitative polymerase chain reaction analysis.

| Target | Forward (5' → 3') | Reverse (5' → 3') |
| --- | --- | --- |
| RPL22 | CACGAAGGAGGAGTGACTGG | TGTGGCACACCACTGACATT |
| GAPDH | AGAACATCATCCCTGCCTCT | ACCCTGTTGCTGTAGCCAAA |
| HPRT1 | CCTGGCGTCGTGATTAGTGAT | AGACGTTCAGTCCTGTCCATAA |
| GFAP | AGAGAGGTCAAGCCCAGGAG | GGTCACCCACAACCCCTACT |
| SERPINA3 | GGTGAGCTCTACCTGCCAAA | CACAGCCTTATGGACCACCT |
| AMIGO2 | GATACTGCAGGGCAGAA | GACGCCACAAAAGGTGTGTC |
| C3 | GAGCCAGGAGTGGACTATGTGTA | CAATGGCCATGATGTACTCG |
| C1QA | GAGCATCCAGTTGGAGTTGAC | ACACAGAGCACCAGCCAT |
| IL-8/CXCL8 | CCCAACCCCTACCTTCTCTC | GTCCACTCTCAATCACTCTCAG |
| CCL22 | GCCTACTCTGATGACCGTGG | AGAGAGTTGGCACAGGCTTC |
| CCL2 | CCAAGCAGAAGTGGGTTCAG | TAAAACAGGGTGTCTGGGGA |
| TNF | TGCACTTTGGAGTGATCGGC | ACAACATGGGCTACAGGCTT |
| IL-6 | AGTCCTGATCCAGTTCCTGC | CTGGCATTTGTGGTTGGGTC |
| AIF-1 | TCATGTCCCTGAAACGAATG | CCAGCATCATCCTGAGAAAG |
| TUBB3 | GGGCCTITGGACATCTCTTC | CCTCCGTGTAGTGACCCTTG |

**Table S2.**

Literature-curated set of relevant cell type-specific marker genes.

| Cell type | Abbreviation | Gene list |
| --- | --- | --- |
| Immature astrocytes | immat-Astro | GFAP, GLUL, NES, VIM |
| Mature astrocytes | m-Astro | GLUL, S100B, SLC1A3, SLC1A2 |
| GABAergic neurons | gaba-Neuron | GAD1, GAD2 |
| Glutamatergic neurons | glu-neuron | GRIN1, GRIN2B, GLUL, GRIN2A |
| Immature neurons | immat-Neuron | NEUROD1,TBR1, TUBB3 |
| Mature neurons | m-Neuron | RBFOX3, MAP2, SYP, MAPT |
| Microglial cells | MGL | P2RY12, P2RY13, CD68, AIF1 |
| Oligodendrocyte precursor cells | OPC | LHFPL3, MEGF11, PCDH15, PDGFRA |
| Oligodendrocytes | m-OL | MBP, MOG,MAG |

**Table S3.**

rAAV serotype 9 vector characteristics.

| Nomenclature | Transgene | Promoter | | Size (bp) | Putative CpG islands | Full capsids (%) |
| --- | --- | --- | --- | --- | --- | --- |
| rAAV9-eGFP(1) | eGFP | CMV promoter | | 2778 | 4 | ~ 50% |
| rAAV9-eGFP(2) |  | CBA promoter | CMV promoter | 4165 | 8 | ~ 83% |
| rAAV9-mCherry | mCherry | CAG enhancer | CAG promoter | 4564 | 6 | ~ 67% |

**Table S4.**

Sequences of SABER-FISH probes for AAV9-mCherry, catalytic hairpin, and fluorescent oligonucleotide (Fluor Oligo).

| ID | Sequence (5' → 3') |
| --- | --- |
| Probe-1 | GGCGGTCAGCCAGGCGGGCCATTTACCGTAAGTTATGTTTCATCATCAT |
| Probe-2 | GACGTCAATGGAAAGTCCCTATTGGCGTTACTATGGGAACTTTCATCATCAT |
| Probe-3 | GGCGTACTTGGCATATGATACACTTGATGTACTGCCAAGTGGGCAGTTTCATCATCAT |
| Probe-4 | GCATAATGCCAGGCGGGCCATTTACCGTCATTGACGTTTCATCATCAT |
| Probe-5 | ACTGCCAAGTAGGAAAGTCCCATAAGGTCATGTACTGTTTCATCATCAT |
| Probe-6 | ATCTCGAACTCGTGGCCGTTCACGGAGCCCTCCATGTGTTTCATCATCAT |
| Probe-7 | GGGTGCTTCACGTAGGCCTTGGAGCCGTACATGAACTTTCATCATCAT |
| Probe-8 | CGCCGTCCTCGAAGTTCATCACGCGCTCCCACTTGAAGTTTCATCATCAT |
| Probe-9 | GGGAAGTTGGTGCCGCGCAGCTTCACCTTGTAGATGAACTTTCATCATCAT |
| Probe-10 | AGTGGCCGCCGTCCTTCAGCTTCAGCCTCTGCTTGATCTTTCATCATCAT |
| Probe-11 | CAGCTGCACGGGCTTCTTGGCCTTGTAGGTGGTCTTGTTTCATCATCAT |
| Probe-12 | CGTTGTGGGAGGTGATGTCCAACTTGATGTTGACGTTGTTTCATCATCAT |
| Probe-13 | CCCTCGGCGCGTTCGTACTGTTCCACGATGGTGTAGTCTTTCATCATCAT |
| Probe-14 | ATGATGGCCATGTTATCCTCCTCGCCCTTGCTCACCATGGTTTCATCATCAT |
| Probe-15 | AATGATGAGACAGCACAATAACCAGCACGTTGCCCAGGAGCTGTTTCATCATCAT |
| Probe-16 | CATGAACATGGTTAGCAGAGGCTCTAGAGCCGCCGGTCACTTTCATCATCAT |
| Probe-17 | GCCTCCCAGATTTCGGCTCCGCACAGATTTGGGACAAAGTTTCATCATCAT |
| Probe-18 | AACAAGCCGTCATTAAACCAAGCGCTAATTACAGCCCGGAGGTTTCATCATCAT |
| Probe-19 | AAAGGAAACTTTCGGAGCGCGCCGCTCTGATTGGCTGTTTCATCATCAT |
| Probe-20 | GGGCTCACCTCGACCATGGTAATAGCGATGACTAATACGTTTCATCATCAT |
| Hairpin | ACATCATCATGGGCCTTTTGGCCCATGATGATGTATGATGATGTTTTTTT |
| Fluor oligo | /5ATTO488N/TTATGATGATGTATGATGATGT |
